# Cerebellar involvement in learning to balance a cart-pole system

**DOI:** 10.1101/586990

**Authors:** Nicolas Ludolph, Thomas M. Ernst, Oliver M. Mueller, Sophia L. Goericke, Martin A. Giese, Dagmar Timmann, Winfried Ilg

## Abstract

The role of the cerebellum in error-based motor adaptation is well examined. In contrast, the involvement of the cerebellum in reward-based motor learning is less clear. In this study, we examined cerebellar involvement in a reward-based motor learning task, namely learning to control a virtual cart-pole system, over five consecutive days. Subjects with focal cerebellar lesions were compared to age-matched controls in terms of learning performance and underlying control mechanisms.

Based on the overall balancing performance we have identified two subgroups of patients: (1) patients with learning performance comparable to healthy controls and (2) patients with decelerated learning, unsaturated learning progress after five days and decreased inter-manual transfer. Furthermore, we found that online learning is impaired while offline learning is partly preserved in cerebellar subjects. Regarding control mechanisms, decreased control performance was associated with impairments in predictive action timing.

Voxel-wise lesion symptom mapping based on the two subgroups revealed strong associations between impairments in controlling the virtual cart-pole system and lesions in intermediate and lateral parts of lobules V and VI. These results together with previous reports suggest that the ability to predict the dynamics of the cart-pole system is an important factor for the reward-based skill acquisition process.

## INTRODUCTION

A large number of studies have shown that cerebellar patients are impaired in motor adaptation by exploiting diverse experimental paradigms, which are based on visuomotor rotation^2–4^, force-field adaptation^5,6^ or split-belt treadmill adaptation^7^ (see reviews by Bastian et al.^8,9^). It is the current opinion that the underlying learning mechanism, which drives motor adaptation and which is impaired in cerebellar patients, is error-based learning of internal forward models. Internal forward models predict the sensor consequences of actions^10^ and are thus suggested to play a crucial role for the predictive control mechanisms in the skilled execution of multi-joint movements^11,12^ as well as for manipulating objects^11,13,14^.

On the other hand, there is increasing evidence, collected by motor training studies, showing that patients with degenerative disease can improve their motor capabilities by intensive motor training^15–17^. The learning mechanisms underlying these improvements in motor control are largely unclear^18,19^.

Because cerebellar patients are impaired in establishing and recalibrating internal forward models using error-based learning, it was hypothesized that cerebellar patients potentially use alternative motor learning mechanisms like reward-based learning or use-dependent learning, which have been suggested to be less dependent on (or even independent from) the integrity of the cerebellum^20,21^.

First evidence in this direction is provided by a recent study on reward-based learning of a visuomotor adaptation paradigm^22^. Although the disease-induced increase in motor variability shown by the cerebellar patients influenced the reward-based learning processes negatively, cerebellar patients were able to learn the altered sensorimotor mapping under a closed-loop reinforcement schedule and retained much more of the learned reaching pattern compared to when they had to perform error-based learning.

In order to explore the preserved capability of reward-based learning in patients with cerebellar dysfunctions further, we here examined cerebellar patients and healthy controls in a reward-based skill acquisition task, namely the skill of balancing a virtual cart-pole system. Cart-pole balancing has been studied in the context of reinforcement learning as a benchmark for computational algorithms^23^ and in the context of exploiting internal forward models for the prediction and control of object dynamics^24–26^.

In a recent study, we have examined the skill acquisition processes in young healthy subjects while learning to balance a cart-pole system in virtual reality^27^. The complexity of the motor task was gradually increased during the learning process by increasing the virtual gravity. Analyses of the task performance and action timing throughout the learning process revealed that the gradual increase of the virtual gravity (1) lead to faster learning but (2) required sensorimotor adaptation of predictive models of the cart-pole system. Furthermore, the results suggested that (3) predictive control mechanisms are crucial to master the cart-pole balancing task.

Exploiting this paradigm of virtual cart-pole balancing, the aim of this study was to examine the involvement of the cerebellum in the skill acquisition process of controlling a complex dynamic system. Specifically, we examined the task performance and action timing of nineteen subjects with focal cerebellar lesions and very mild to moderate ataxia symptoms in short sessions over five subsequent days. Examining the learning processes over several days enabled us to examine online as well as offline learning^28,29^. In addition to the analysis of behaviour, we used voxel-wise lesion symptom mapping (VLSM) to associate specific behavioural impairments with focal cerebellar damage in order to elucidate which cerebellar regions play a functional role in cart-pole balancing.

Our main hypothesis was that even focal lesions, which result only in very mild to moderate ataxia symptoms, might lead to recognisable changes in learning performance. Furthermore, due to the potential involvement of the cerebellum in the acquisition of the cart-pole skill, we hypothesized different areas of the cerebellum as candidates, in which lesions could be associated with impaired learning capacity. Namely,

i. We hypothesized that cerebellar damage in areas representing the online state estimator^30^ for the control of arm and hand movements will lead to increased motor variability in space and time, which likely effects also the reward-learning process^22^.
ii. In addition, lesions in areas representing forward models of the cart-pole system could also influence the skill acquisition process. More posterior and lateral regions of the cerebellum could represent the dynamic model of the cart-pole system^31^.
iii. The deep cerebellar nuclei, in particular the interposed and dentate nuclei, as the output of the relevant cerebellar areas, could also influence the sensorimotor execution and learning behaviour.

## METHODS

### Subjects

Nineteen patients (mean age 25.6 years, 12 female, 7 male) with chronic surgical cerebellar lesions (**Table 1**) and nineteen age-matched self-reportedly neurologically healthy control subjects participated in the experiment. Twelve patients suffered from pilocytic astrocytoma (that is, astrocytoma WHO grade I), two from astrocytoma WHO grade II, four from vascular tumors and one from a dermoid cyst. None of the cerebellar subjects had received adjuvant chemo-or radiotherapy. Patients showed very mild to moderate signs of ataxia as examined by an experienced neurologist (D. T.). Severity of ataxia was rated using the International Cooperative Ataxia Rating Scale ICARS (Range: 0-20, Mean: 4.68 (maximum possible ICARS score = 100), see **Table 1**, Trouillas et al.^1^). Handedness was determined using the Edinburgh Handedness Inventory^32^.

**Table 1.** List of the examined young adult subjects with chronic surgical cerebellar lesions. We decided to let subjects who had to switch handedness after their surgery (cp7 and cp12) train the cart-pole balancing with their “new dominant” hand as indicated in the table. Listed are the total and ICARS ^1^=^1^ subscores (P) Posture and Gait, (K) Kinetic function, (S) Speech, and (O) Oculomotor function. For cerebellar subject 8 (marked by *) we were not able to measure the fifth day due to a technical error.

| ID | Age (yr) | Gender | Handedness (after surgery) | Diagnosis / Primary localization | Time (yr) since surgery | ICARS |  |  |  |  |
| --- | --- | --- | --- | --- | --- | --- | --- | --- | --- | --- |
|  |  |  |  |  |  | P (max. 34) | K (max. 52) | S (max. 6) | O (max. 8) | Total (max. 100) |
| cp01 | 32 | M | R | Hemangioblastoma (L) | 7.25 | 0 | 0 | 0 | 0 | 0 |
| cp02 | 22 | F | L | Pilocytic astrocytoma (mid', R) | 17.5 | 2.5 | 2 | 0 | 0 | 4.5 |
| cp03 | 25 | M | R | Angioma (L) | 5.5 | 3.5 | 11 | 0.5 | 5 | 20 |
| cp04 | 40 | F | R | Pilocytic astrocytoma (R) | 8.75 | 0 | 0 | 0 | 0 | 0 |
| cp05 | 28 | F | R (L) | Pilocytic astrocytoma (R) | 18.25 | 2 | 6 | 0 | 6 | 14 |
| cp06 | 27 | F | R | Pilocytic astrocytoma (L) | 12 | 0 | 0 | 0 | 0 | 0 |
| cp07 | 24 | F | R | Pilocytic astrocytoma (mid, L) | 8.25 | 2 | 3.5 | 0 | 5 | 10.5 |
| cp08* | 30 | M | R | Astrocytoma WHO Grade 2 (mid-L) | 14.5 | 2 | 0 | 0 | 0 | 2 |
| cp09 | 27 | M | R | Astrocytoma WHO Grade 2 (L, mid) | 12.75 | 0 | 0 | 0 | 3.5 | 3.5 |
| cp10 | 27 | F | R | Pilocytic astrocytoma (mid) | 18.5 | 2 | 0 | 0 | 0 | 2 |
| cp11 | 36 | F | R | Hemangioblastoma (L) | 10.5 | 0 | 0 | 0 | 0 | 0 |
| cp12 | 23 | F | R (L) | Pilocytic astrocytoma (R) | 20 | 5 | 6 | 0 | 6 | 17 |
| cp13 | 29 | F | R | Pilocytic astrocytoma (mid) | 20 | 2.5 | 2 | 0 | 0 | 4.5 |
| cp14 | 19 | M | R | Pilocytic astrocytoma (L) | 11.75 | 0 | 0 | 0 | 0 | 0 |
| cp15 | 19 | F | R | Dermoid cyst (mid) | 8.75 | 0 | 0 | 0 | 0 | 0 |
| cp16 | 24 | F | R | Pilocytic astrocytoma (L) | 9 | 2 | 0 | 0 | 6 | 8 |
| cp17 | 17 | M | R | Pilocytic astrocytoma (R) | 10 | 0 | 0 | 0 | 0 | 0 |
| cp18 | 19 | M | R | Cavernoma (R) | 1 | 0 | 0 | 0 | 0 | 0 |
| cp19 | 18 | F | R | Pilocytic astrocytoma (R) | 1.5 | 2 | 1 | 0 | 0 | 3 |

The study was carried out according to standard guidelines and regulations and was approved by the ethical review board of the medical faculty of the Eberhard-Karls-University and university clinics in Tuebingen as well as by the ethical review board of the Essen University Hospital (Tuebingen 409/2014BO2, Essen 14-6053-BO). All participants, and in one case due to minority the legal guardians, gave informed written consent prior to participation.

### Behavioural Experiment

#### Cart-Pole Balancing Task

In the *cart-pole balancing task*, subjects have to learn to control a virtual simulation of the cart-pole system (**Fig 1** A). Gravity forces the pole to rotate downwards when not being perfectly upright. The goal of the cart-pole balancing task is to keep the pole upright. Specifically, the pole has to remain within the green circular segment (±60 degree, **Fig 1** A) while the cart must not leave the track (±5 m). By repeatedly accelerating the cart at the correct time to the left or right depending on the pole angle, it is possible to prevent the pole from falling. Participants accelerated the cart by applying virtual forces using a haptic input device (3Dconnexion SpaceMouse® Pro, **Fig 1** B, for technical details see Supplementary Material S1). A trial was considered successful if balance was maintained for 30 seconds without violating any of the two given constraints.

**Fig 1.**
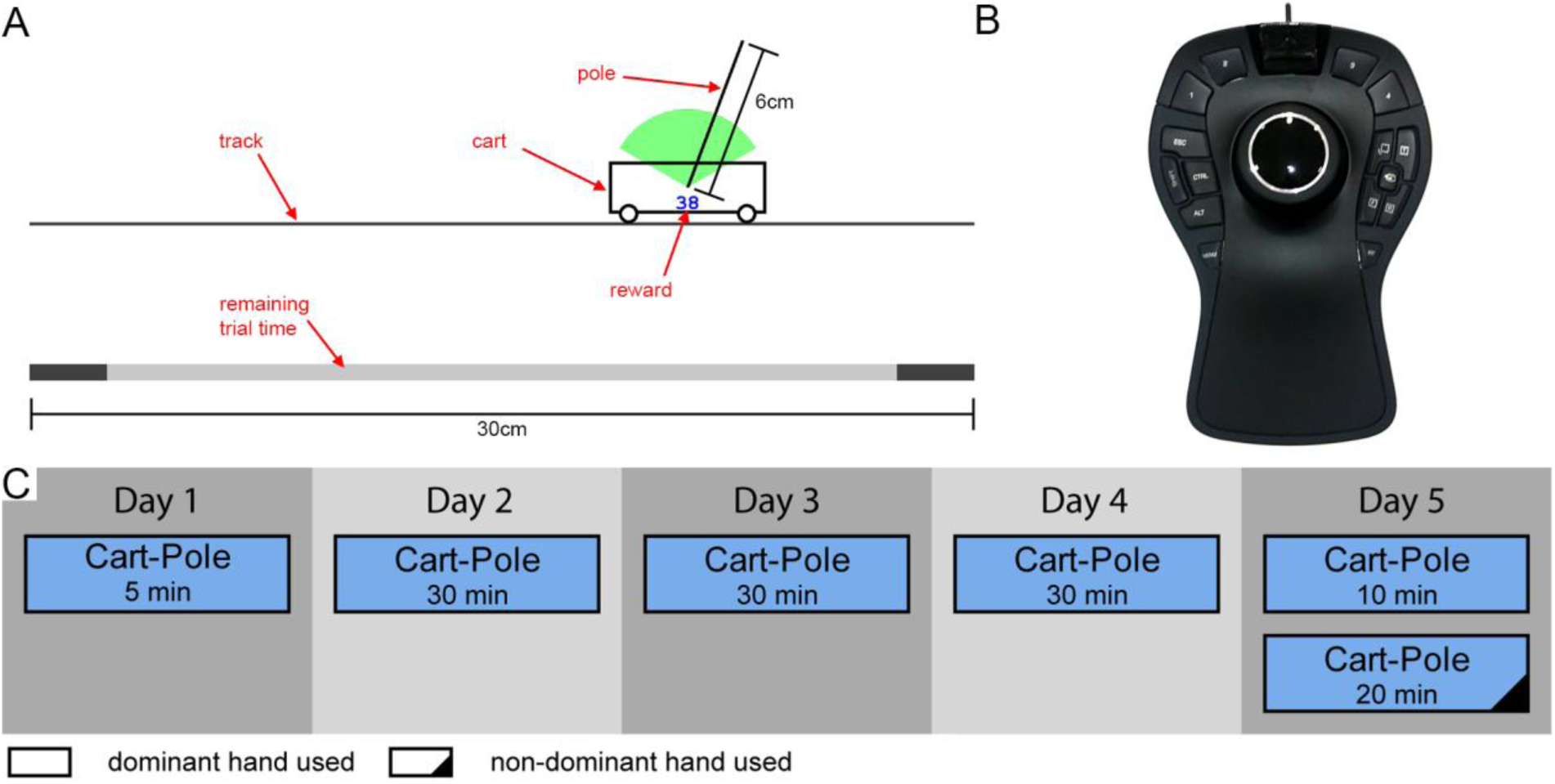
Cart-Pole balancing task, input device and experimental protocol. (A) In the cart-pole balancing task, the pole has to be balanced on the cart by accelerating the cart to the left and right. On the bottom, the remaining trial time is indicated. The two dark areas meet in the centre after 30 seconds. (B) The left-right translation degree-of-freedom of the input device was used to control the virtual force, which in turn accelerated the cart. (C) The duration and hand used for each of the training session on the five consecutive days of the experiment.

In order to reduce the initial difficulty, the task started with the gravitational constant set to g_init_=1.0m/s^2^. After every successful trial, the gravity was increased by 0.1m/s^2^ until the maximum of g_max_=3.0m/s^2^ was reached. Thus, due to this performance-dependent increase, every subject was exposed to an individual gravity profile over the course of the experiment. Like in our previous study^27^, we displayed a cumulative reward as number within the cart, which increases over the duration of the trial depending on the state of the system and the applied virtual force. Herewith, we gave skilled subjects the opportunity to improve further when having reached the maximum gravity already and being able to balance the system reliably for 30 seconds.

#### Experimental Protocol

Overall, the experimental protocol involved five consecutive days (**Fig 1** C). On the first day, we let subjects get a first impression (5 minutes) of the cart-pole balancing task. We made sure that all participants were sufficiently familiar with the experimental setup itself in order to qualify them to perform the remaining sessions without our supervision at home. The subsequent three days (day 2-4) constitute the main cart-pole balancing training, lasting 30 minutes per session. After the extensive cart-pole balancing training over three days, the last day (day 5) was devoted to examining the transfer to the non-dominant hand. On this last day, participants first again performed the cart-pole balancing task using their dominant hand for 10 minutes and then switched to their non-dominant hand to perform the task for another 20 minutes.

#### Handedness and lesion side

Focal cerebellar damage affects primarily the control of the ipsilateral limb. Consequently, previous studies have tested the ipsilateral hand to the lesion independent of handedness (e.g. Donchin et al.^2^) to examine the consequences of cerebellar damage on the motor performance. Learning complex tasks with the non-dominant hand can however be extremely difficult especially if fine motor control is required like in the cart-pole balancing task. Moreover, according to Schlerf et al.^33^, both cerebellar hemispheres are active during complex manual tasks. This suggests that the performance in complex tasks, such as the cart-pole balancing task, is affected independent of lesion side. Thus, we decided that all subjects acquire the cart-pole balancing task using their dominant hand, which was not necessarily the ipsilateral hand to the lesion, but we also examined the non-dominant hand after the initial skill acquisition.

### Data Processing and Analysis

#### Measures of task performance and action timing

We analysed the observed behaviour regarding three measures: (1) the maximum reached gravity level (mrGL), (2) the trial length, (3) and the action timing and variability^27^. Like in our previous study^27^, the cumulative reward at the end of the trial was highly correlated with the trial length and did not provide any further insight. An overview of all measures is provided in **Table 2**.

**Table 2.** Overview of the examined measures.

| Measure | Short Description |
| --- | --- |
| **mrGL** | The maximum reached gravity level (mrGL) is the final gravity level that was reached during the training phase. It measures the progression of learning because the gravity was only increased if balance was maintained successfully for 30 seconds. Due to the experimental setup, mrGL is limited by $g_{init}=1.0m/s^2$ and $g_{max}=3.0m/s^2$. |
| **rnTL** | The reciprocal normalized trial length (rnTL) is the reciprocal value of the normalized trial length ($TL/T_0$) which relates the measured trial length (TL) to the hypothetical trial length if no controlling force was applied ($T_0$).<br>• dominant hand Per trial while using the dominant hand (0-105min).<br>• non-dominant hand Per trial while using the non-dominant hand (105-125min).<br>• both hands combined using the averages over 100-105min and 120-125min. |
| **AT** | The action timing (AT) for a specific pole angle event is the average duration between the event occurrences and the respective changes in the applied virtual force direction. We analysed the action timing as average over multiple pole angles.<br>• dominant hand Per 5mins while using the dominant hand (0-105min).<br>• non-dominant hand Per 5mins while using the non-dominant hand (105-125min).<br>• both hands combined using the averages over 100-105min and 120-125min. |
| **AV** | The action variability (AV) is defined by the mean standard deviation of the applied virtual forces in a centred window of 120ms length around the estimated action timing for a specific pole angle event. We analysed the action variability as average over multiple pole angles. |

**(1)** The *maximum reached gravity level* (mrGL) is the coarsest measure of performance in the cart-pole balancing task. Because the gravity was only increased if balance was maintained successfully for 30 seconds, the mrGL measures the progression of learning. The maximum reachable gravity level was set to g_max_=3.0m/s^2^.

**(2)** A more fine-grained measure of performance is the *trial length* (TL). However, due to the individual increase in gravity and the therewith-associated difference in difficulty, it is necessary to normalize the trial length. The normalized trial length (TL/T_0_) relates the measured trial length (TL) to the hypothetical trial length if no controlling force was applied (T_0_). The normalized trial length represents thereby a multiple of being better than doing nothing^27^. For the statistical analysis, we used the *reciprocal of the normalized trial length* (rnTL=T_0_/TL) in order to equalize between-subject variability.

**(3)** The *action timing* (AT) and *action variability* (AV) allow us to quantify the changes in the subjects’ actions over the course of learning. In order to obtain a measure of action timing, we examined the applied virtual forces of each subject as function of the system state, specifically as function of the pole angle (**Fig 2** A)^27^. Calculating this measure involves the definition of events in the state space and analysing the applied force relative to those events (event-triggered averaging, **Fig 2** B). We focused our analysis on the situations when the pole is tilted by a certain angle and is rotating downwards. In these situations, which we describe as events, a counter-action is necessary. We refer to the timing of these counter-actions relative to the previously defined events as action timing. By averaging these time estimates across all events within bins of 5min, we gain a measure of general action timing performance in the cart-pole balancing task. Furthermore, we estimated the variability of the actions by calculating the mean standard deviation of the applied forces in a centred window of 120ms length around the estimated action timing (**Fig 2** C). More computational details of this method can be found in Ludolph et al.^27^.

**Fig 2.**
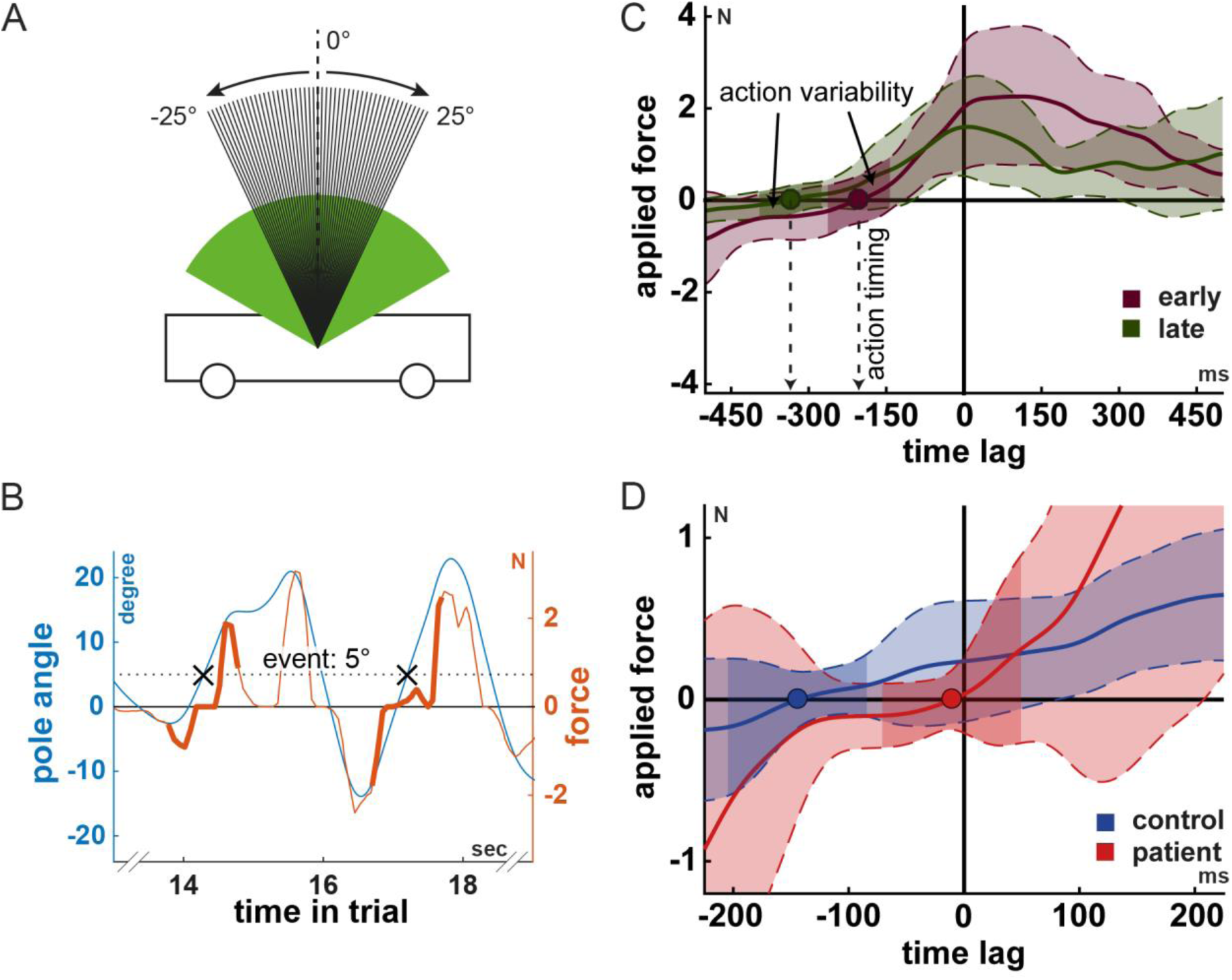
Action timing and action variability in the cart-pole balancing task. (A) All pole angles investigated as events (integer valued pole angles from −25° to 25°). The arrows indicate the direction of the pole movement. (B) Pole angle (blue) and input force (orange) trajectories illustrating two event occurrences (black crosses) and corresponding two force segments (think lines). (C) Average force segments for two periods (early vs. late) during learning for illustrating the action timing and variability measures. The dots indicate the action timing. Negative and positive time lags represent the time before and after the event, which has zero time lag. The dark coloured areas illustrate the action variability measure. The lightly shaded areas express the overall variability in force segments. The panels (A-C) have been reused with minor modifications from Ludolph et al.^26^. (D) Comparison of the action timing between a control and a cerebellar patient for the gravity of g=2.6m/s^2^. The action timing was determined over the last few trials on that gravity level for both subjects. While the control subject mastered that gravity level, the cerebellar subject failed to balance the cart-pole system on that gravity level. Notice that the action timing (red dot) of the patient is considerably less predictive than the action timing of the control subject (blue dot).

### Analysis of task performance and action timing across hands

In order to examine the inter-manual transfer of learned motor behaviour we analysed the change in task performance (rnTL) from the last session with the dominant hand to the session in which subjects used their non-dominant hand. Hereto, we subtracted the average task performance measured during the two periods (100-105min vs 105-110min) for each subject separately and performed a group comparison. Furthermore, we examined the relation of the estimated action timing between both hands. Our hypothesis was that if the representation of predicting the behaviour of the cart-pole system is hand independent from the hand trained with there should be a strong correlation between the estimated action timing for each hand.

### Combining the performance of both hands

In addition to the task performance (rnTL) and action timing (AT) of each hand, we defined measures that represent the performance of both hands. To this end, we calculated the average of these measures over the last 5 minutes of the two blocks on the fifth day (100-105min and 120-125min). When plotting the task performance (rnTL) of both hands against each other (Supplementary Material S3), the general ability to balance the cart-pole system is represented by the distance from zero (radius in polar coordinates). Similarly, we defined the measure “AT combined hands” in order to capture the general ability to time actions in the cart-pole task. Based on the pairs (dominant, non-dominant hand) of estimated action timings (AT), we calculated the empirical median and median absolute deviation of the control subjects’ performances. Using these estimates, we calculated the modified z-scores^34^ of each pair yielding lower scores for normal (represented by the control group) and higher scores for abnormal behaviour.

### Online and offline learning

Improvement during training sessions is called *online learning*, while improvement between sessions without actual training, is called *offline learning*^28^. In order to evaluate and compare these phases of learning, we averaged the rnTL over the first and last 5 minutes of each session yielding two measures (initial and end performance) per session. We then calculated the *online change in rnTL* by subtracting the initial from the end performance of the same session and the *offline change in rnTL* by subtracting the end performance of the previous session from the initial performance of the subsequent session. We excluded the first day from this investigation because it lasted only five minutes overall.

### Statistical analysis of behavioural data

Statistical analysis has been performed using MATLAB 2016b (The MathWorks, Inc.) and SPSS 23 (IBM Cooperation). Simple group comparisons have been performed using Wilcoxon’s rank sum test or two-sided t-test depending on the result of Kolmogorow-Smirnow-Lilliefors’ test. Repeated measures ANOVA (rm-ANOVA) has been performed to compare group performance across days. Greenhouse-Geisser (GH) and Bonferroni adjustments have been performed where appropriate. Where adequate we expressed corrected p-values as p and uncorrected p-values as p^u^. Bivariate correlations were performed using Spearman’s rank correlation coefficient. In order to compare the decrease in rnTL between groups, we fitted a non-linear regression model (see detail in Supplementary Material S2).

### Lesion symptom analysis

Anatomical magnetic resonance (MR) images of all cerebellar subjects were acquired using a 3-T Siemens scanner (Skyra) with a 32-channel head coil (see for more technical details Supplementary Material S3).

Cerebellar lesions were manually traced on axial, sagittal, and coronal slices of the non-normalized 3-D MPRAGE MRI data set and saved as regions of interest (ROI) using the free MRIcron software (http://www.mricro.com/mricron). Where appropriate ROIs were adjusted based on lesion extent in FLAIR images. Images and ROIs were normalized using a spatially unbiased infratentorial template of the cerebellum (SUIT; Diedrichsen et al.^35^) with the SUIT toolbox in SPM8 (Wellcome Department of Cognitive Neurology, London, UK).

For performing the voxel-wise lesion symptom mapping (VLSM), all lesions were flipped to the right. VLSM was performed with the use of MRIcron and NPM software (included in MRIcron).

Associations between cerebellar damage and behavioural impairments were obtained using subtraction analysis^36^ in combination with statistical confirmation using multiple Liebermeister’s tests^37^. For the Liebermeister’s tests only voxels damaged in at least 16% of individuals (n=3) were considered. As recommended by Rorden et al.^37^, we corrected the significance level using permutation thresholding (4000 permutations). In order to perform these tests, we grouped cerebellar subjects into an affected and unaffected subgroup based on the performance of the control subjects in the measure under investigation.

### Data availability statement

The datasets generated during and/or analysed during the current study are available from the corresponding author on reasonable request.

## RESULTS

### Behavioural analysis of the cart-pole task

#### Maximally reached gravity level is significantly lower for cerebellar patients

Subjects increased the gravity level over the course of learning by balancing the cart-pole system successfully (**Fig 3** A). We noticed that not all subjects reached the maximum gravity level of g_max_=3.0m/s^2^. For each subject, we thus determined the maximally reached gravity level (mrGL) as measure of overall capability to acquire the cart-pole balancing skill. While all control subjects reached the maximum gravity level (g_max_=3.0m/s^2^), only 57% of the cerebellar subjects reached this level (**Fig 3** C). Consequently, the maximum reached gravity level is significantly lower for the group of cerebellar subjects (Wilcoxon’s ranksum test, p<0.01). We categorize subjects, who mastered at least half of the gravity levels (mrGL>g_thresh_=2.0m/s^2^, **Fig 3** D, see Methods), as unaffected and those subjects, who did not master at least half of the gravity levels, as affected (mrGL≤g_thresh_=2.0m/s^2^). This method lead to three groups: (1) affected (N=5, CPa, ICARS range [3-20], mean 11.0), (2) unaffected cerebellar (N=14, CPu, ICARS range [0-14], mean 2.42) and (3) control subjects (N=19, HC). Affected and unaffected patient groups differed significantly from each other in the clinical ataxia score ICARS (p=0.0076). However, also in the unaffected cerebellar group there were five subjects with mild to moderate ataxia symptoms (ICARS range [2-14], mean 5.1).

**Fig 3.**
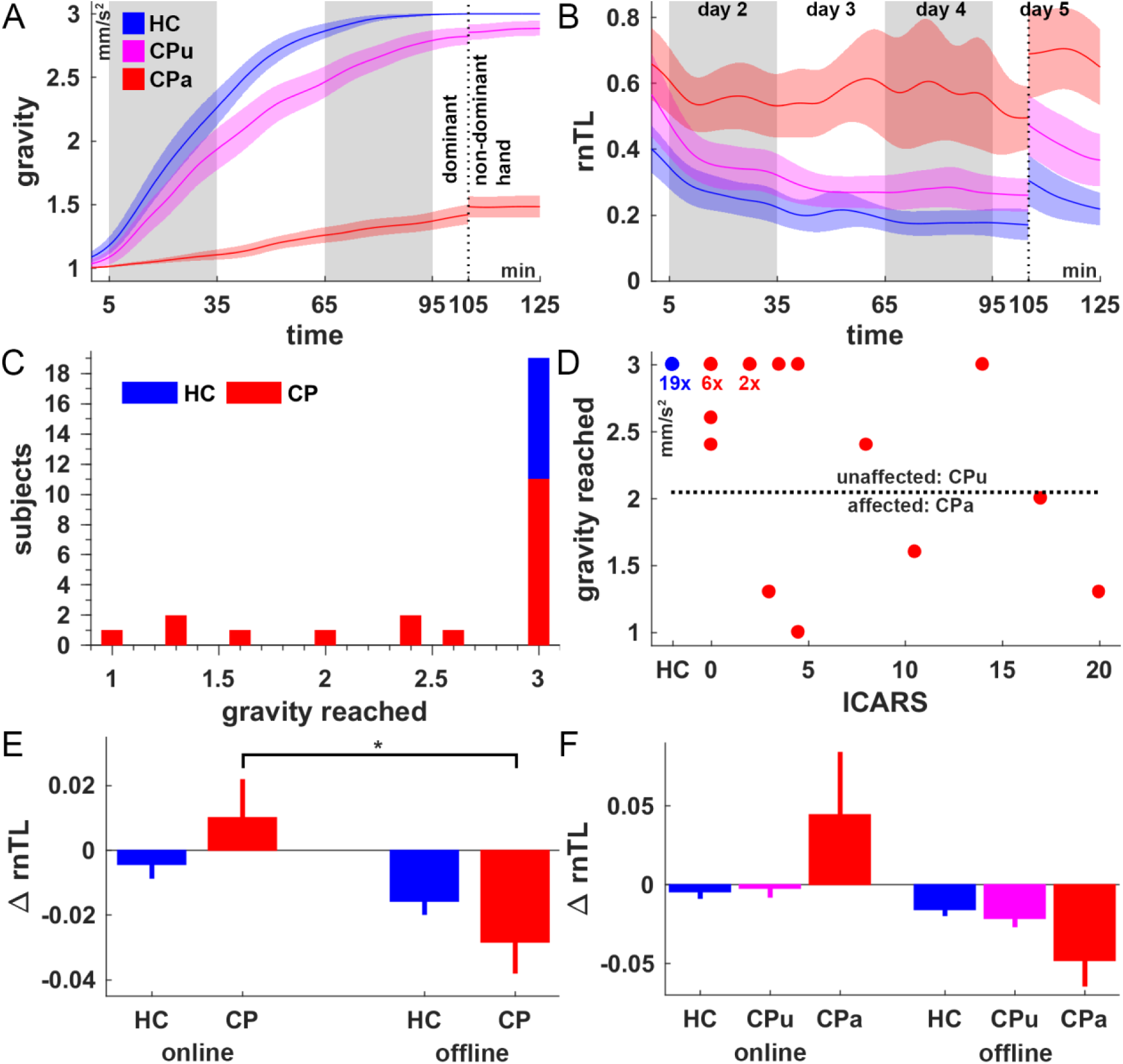
Performance in the cart-pole balancing task. (A) Average gravity and (B) reciprocal normalized trial length (rnTL) across time for affected (CPa, red), unaffected cerebellar (CPu, magenta) and healthy control subjects (HC, blue). Good performance is associated with low rnTL. Cerebellar subjects were classified as unaffected if they reached a higher gravity than g_thresh_=2.0m/s^2^. Dashed lines indicate the switch from dominant to non-dominant hand usage. For visualization purposes, the curves were smoothed over time using the weighted running average method. (C) Frequency of the reached gravity over all sessions for cerebellar subjects (red) and control subjects (blue). (D) Maximum reached gravity level (mrGL) as function of the ICARS score for cerebellar (red) and control subjects (blue). The dotted line represents the thresholds of impaired behaviour and splits the cerebellar subjects into an affected (CPa) and unaffected (CPu) subgroup. (E, F) Online and offline learning measured by the change in performance (rnTL) within sessions (online) and between sessions (offline) for cerebellar (red) and control subjects (blue), and in (F) also for the two cerebellar subgroups. Error bars indicate ±1 S.E.M. (* p<0.05)

#### Learning is significantly slowed down for affected cerebellar patients

The reciprocal normalized trial length (rnTL) resolves the task performance more fine grained (see Methods). Both groups, patients and controls, decreased the reciprocal normalized trial length significantly over the course of training (Spearman’s correlation coefficient, time vs. rnTL, cerebellar: r=-0.25, p<0.001; control subjects: r=-0.48, p<0.001) (**Fig 3** B). By fitting a non-linear regression model (see Supplementary Material S2), we found significantly different rates of learning between affected and unaffected cerebellar subjects (F=23.52, p<0.001, Bonferroni corrected) as well as between affected cerebellar and control subjects (F=25.19, p<0.001, Bonferroni corrected). Between unaffected cerebellar and control subjects we did not find any significant difference in learning rate (F=1.07, p^u^=0.34).

#### Affected cerebellar patients are impaired in online but show offline learning

We analysed the relationship between online and offline learning (see Methods) based on the main training phase (days 2, 3 and 4, **Fig 3** E). Rm-ANOVA revealed a significant effect of learning phase (F=4.758, p=0.036, GH) on the task performance (rnTL), while the factors day and group did not reach significance (day: F=0.839, p=0.40, GH; group: F=0.185, p=0.67). None of the interactions reached significance. Post-hoc test revealed that the improvement of cerebellar patients during the offline phases is significantly higher than during the online phases (mean online: 0.01, offline: −0.03, p=0.02, Bonferroni corrected). For control subjects, there was no significant difference between learning phases (mean online: −0.01, offline: −0.02, p=0.48, Bonferroni corrected). Notice that cerebellar subjects even tend to get worse during the practice sessions while getting better across sessions (positive ΔrnLT, **Fig 3** E) suggesting that online learning is impaired while offline learning is intact.

We performed this analysis also using the two subgroups of cerebellar subjects (**Fig 3** F). Although **Fig 3** F suggests that only affected cerebellar patients (mrGL ≤ 2.0m/s^2^) show abnormal online learning behaviour, the small group size of the affected cerebellar subjects did not allow performing an ANOVA.

#### Cerebellar patients show increased action variability and impaired predictive action timing

We found that subjects in the affected cerebellar group show an increased action variability (AV) at the end of the training (t-test, affected cerebellar vs. control: p^u^=0.025, p=0.075; affected vs. unaffected cerebellar: p^u^=0.035, p=0.105) while there was no difference between the unaffected cerebellar and control subjects (t-test, p=0.77).

Analysis of the action timing over the course of the experiment (AT, **Fig 4** A) revealed a significant decrease for the control group (p<0.001) and unaffected cerebellar subgroup (p=0.03) but not for affected cerebellar subjects (p=0.09, see Supplementary Material S2). Notice, that a negative action timing is associated with performing the actions predictively ^27^. There was no significant difference in the action timing learning rate (see Methods) between the unaffected cerebellar and control group (F=1.2422, p^u^=0.27, uncorrected) but affected cerebellar subject showed significantly different (lower) learning rates (vs. control: F=12.0570, p=0.002; vs. unaffected cerebellar: F=6.6918, p=0.03; both Bonferroni corrected).

**Fig 4.**
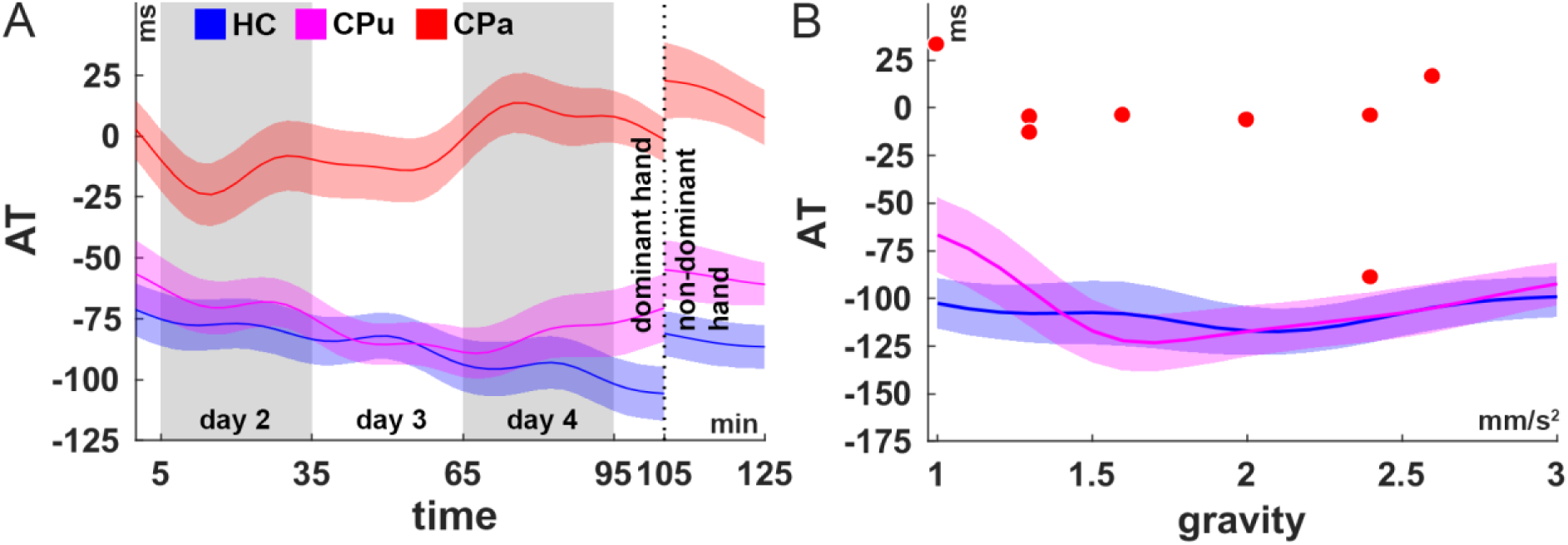
Action timing in the cart-pole balancing task. (A) Action timing (AT) across time for affected (red), unaffected (magenta) cerebellar and control (blue) subjects. For visualization purposes, the data was smoothed in time and the shaded areas indicate ±0.25 SD. (B) Action timing (AT) across the different gravity levels. The action timing of cerebellar subjects, who did not reach the maximum gravity level (red dots), is only shown on the individual maximum reached gravity level as dot. For unaffected cerebellar (magenta) and control (blue) subjects, the action timing over the gravity levels is smoothed for visualization purposes and the shaded areas indicate ±0.25 SD. Notice that the action timing of the cerebellar subjects, who did not reach the maximum gravity (red dots), is much higher (less predictive) than for the control subjects (compare also to **Fig 2**D).

In order to determine the value of predictive action timing regarding mastering the next gravity step, we examined the action timing on the maximum reached gravity level of all cerebellar subjects, who did not reach the maximum gravity level of g_max_=3.0m/s^2^. We found that the action timing for these subjects is significantly less predictive than the average action timing of the control subjects (**Fig 4** B, p<0.01, N=8), indicating that, for mastering the next gravity step, it is necessary to time the actions more predictively than these cerebellar subjects were able to do.

#### Cerebellar patients are impaired in intermanual transfer

When switching from the dominant to the non-dominant hand on the 5th day, performance was significantly more reduced (t-test, p=0.019) for the cerebellar group (N=18, mean change in rnTL: +0.13) compared to healthy controls (N=19, mean change in rnTL: +0.055). This suggests that the transfer from the dominant to the non-dominant hand is less complete in cerebellar patients than in control subjects (**Fig** 3 B, Supplementary Material S3).

#### Action timing for the dominant and non-dominant hand is significantly correlated

In order to examine the potential learning of hand independent representations, we analysed the action timing expressed by both hands. The action timing (**Fig 5** A) is significantly correlated between hands for both groups: healthy controls (Spearman’s correlation coefficient, rho=0.74, p<0.001, N=19) and cerebellar subjects (Spearman’s correlation coefficient, rho=0.54, p=0.02, N=18). Suggesting that the ability to time the actions correctly is rapidly transferred from the training to the non-training hand.

**Fig 5.**
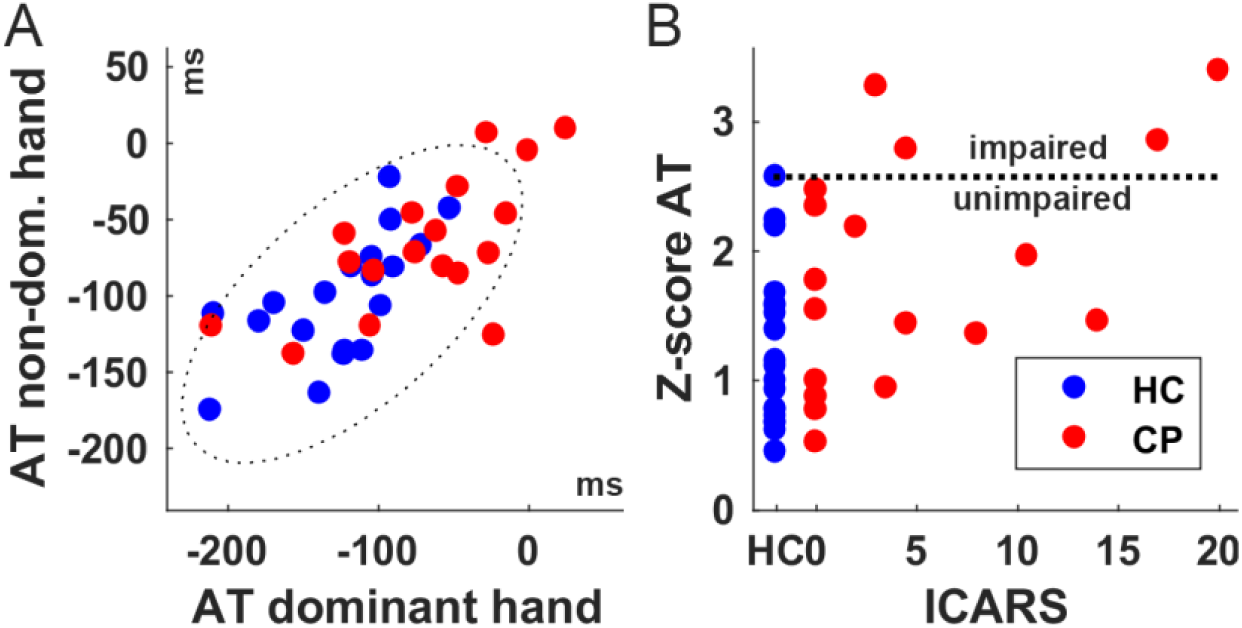
Classification of the action timing behaviour. (A) AT of the dominant versus the non-dominant hand over the last 5 minutes of the respective sessions on the fifth day. The dotted ellipse represents the threshold of being affected. (B) Modified z-score of the action timing (AT combined hands) as function of the ICARS score for cerebellar (red) and control subjects (blue). Being affected in this measure corresponds to the iCPB classification.

### Lesion symptom analysis

#### Lesions are distributed across the cerebellum with focus on the midline and intermediate parts

In twelve cerebellar subjects, lesions involved primarily the cerebellar hemispheres and, in the seven subjects, primarily the midline. Superposition of individual lesions shows that the lesions are distributed across the whole cerebellum with a focus on the midline and intermediate parts (**Fig 6** A). Maxima of overall lesion overlap were found in lobule V, VIIIa, IX (N=8), as well as in lobule VI, VIIb, Crus I and Crus II (N=9). In ten subjects the dentate, and in eight subjects the interposed nuclei were at least partially damaged (maximum lesion overlap: N=7 and N=6, respectively).

**Fig 6.**
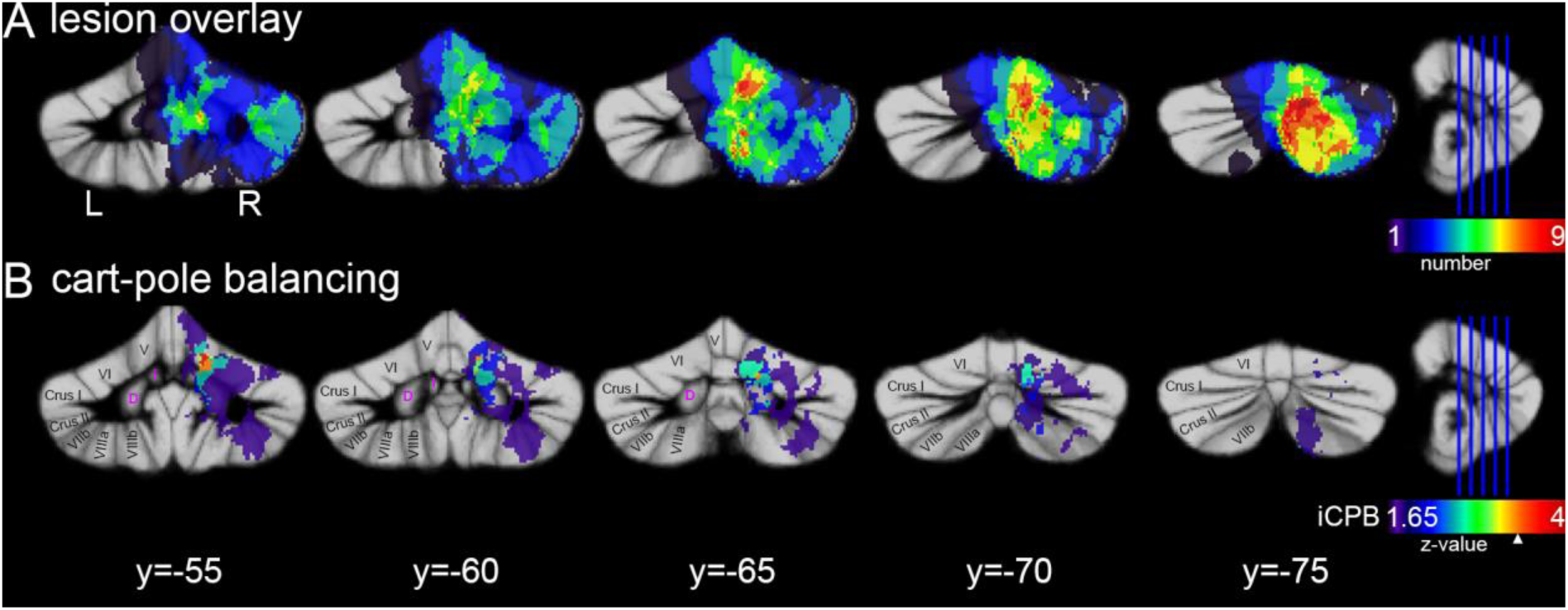
Voxel-wise lesion symptom mapping. (A) Superposition of cerebellar lesions across all cerebellar subjects (N=19) after flipping to the right. Colour indicates the number lesions, which overlap in the given voxel. (B) VLSM considering the general performance in the cart-pole balancing task (iCPB, i.e. impaired according to the measures mrGL, rnTL combined hands and AT combined hands). Shown is the Z-value map of the performed Liebermeister test. The z-value corresponding to the permutation corrected 5% significance level is indicated by a white triangle. All results are superimposed on the maximum probability SUIT template of the cerebellum^71^. Names of cerebellar lobules are indicated according to Schmahmann et al.^72^.

#### Cerebellar regions associated with cart-pole balancing

Based on the task performance and action timing measures, overall six cerebellar subjects were classified as impaired. Of these six subjects, all were classified impaired regarding the task performance (rnTL combined hands, Supplementary Material S3), five were classified impaired because of the low maximum reached gravity level (mrGL≤g_thresh_=2.0m/s^2^, **Fig 3** D) and four were additionally impaired in timing their actions in comparison to the control group (AT combined hands, **Fig 5** A, B). In the following, we call the subgroup of four subjects who showed impaired behaviour in all of these measures, i.e. those who showed impaired action timing behaviour, impaired in cart-pole balancing (iCPB, **Fig 5** B). Here we focus on the iCPB classification, while the VLSM for the individual measures and hands is reported as supplementary material (Supplementary Material S3).

Subtraction analysis based on the general cart-pole balancing performance classification (iCPB, **Fig 5** B, **Fig 6** B) revealed highest consistency in lobule V (100%), VI (92%) and in the interposed nucleus (92%). Slightly lower consistency was found in the dentate nucleus (67%), lobule VIIb (60.7%) and VIIIa (50%). Statistical testing using multiple Liebermeister tests and correction for multiple comparisons (criterion: p<0.05, permutation corrected: Z>3.36) confirmed the clusters in lobule V (max: x=13, y=-56, z=-20), VI (max: x=14, y=-64, z=-26) and interposed nucleus (max: x=11, y=-57, z=-26).

## DISCUSSION

In this study, we have investigated the influence of focal cerebellar lesions on a reward-based motor skill acquisition task, namely learning to control a virtual cart-pole system.

Over a period of 5 days, one subgroup of patients showed learning capabilities comparable with healthy controls, although some of them also showed mild to moderate ataxia signs in the clinical ataxia score (ICARS [0-14]). The subgroup of patients affected in virtual cart-pole balancing (ICARS range [3-20]) showed decreased learning capabilities as well as impaired transfer of control knowledge to the other hand. Comparison of the two subgroups, using lesion symptom mapping (LSM), revealed distinct clusters of voxels associated with impairments in the intermediate and lateral parts of lobules V and VI as well as in interposed and dentate nuclei.

### Behavioural analysis of the cart-pole task

The group of affected patients had a significant higher clinical ICARS score as the unaffected group (p=0.0076). Nevertheless, there were also five patients in the unaffected group (N=14) showing mild to moderate ataxia signs (ICARS range [2-14]). Thus, the clinical score alone cannot explain the behavioural differences in cart-pole balancing.

#### Affected cerebellar patients are impaired in online but show offline learning

Interestingly, we have observed that cerebellar subjects improve significantly more between practice sessions (offline learning) compared to their improvement during the practice sessions (online learning), while there was no such difference for control subjects.

It has been argued that online and offline learning mechanisms might rely on the activity of different brain areas (see Dayan et al.^28^ for a recent review). Studies examining the non-invasive stimulation of different areas of the motor network have shown that stimulating the cerebellum can lead to faster online learning whereas stimulating the primary motor cortex (M1) can facilitate retention and offline learning^29,38,39^. There are however almost no systematic studies on motor learning in cerebellar patients over several days. One of the few studies^40^ showed consolidation of timing improvements in an eye-blink conditioning paradigm over multiple days in patients with focal cerebellar lesions. Together, these studies point in a direction, consistent with our finding, which is that specifically online learning is decreased in cerebellar subjects during skill acquisition while offline learning as well as retention are at least partly preserved in mildly to moderately affected patients with focal cerebellar lesions.

#### Affected cerebellar patients need potentially longer training durations

Another observation regarding the group of affected patients is that the learning curves are not saturated after the learning period of five days (**Fig 3** A), which strongly indicates that these patients would profit from an even longer period of training. In fact, it has been previously stated that learning processes are slowed down in cerebellar patients and that they might therefore be dependent on even longer training durations^19^. Validating this hypothesis would be of particular relevance for continuous training approaches^18^.

#### Affected cerebellar patients exhibit increased action variability

One of our hypotheses was that cerebellar-induced motor impairments result in increased motor action variability and thereby influence the learning process negatively.

In fact, we found that subjects in the affected cerebellar group show increased action variability (AV) at the end of the training. This result is in correspondence with a recent study on reward-based learning in a visuomotor adaptation paradigm involving cerebellar patients. Therrien et al.^22^ found that learning was dependent on the balance of motor noise and exploration variability, with the patient group having greater motor noise and thus learned less in the same time. The authors concluded that cerebellar damage may indirectly impair reinforcement learning by increasing motor noise, rather than interfering with the reinforcement mechanism itself ^22^. The influence of motor variability on reward-based learning has been shown also in other movement disorders like dystonia^41^, supporting the general hypothesis that reward-based learning is dependent on the balance of motor noise and exploration variability.

#### Predictive action timing is crucial

Another of our central hypotheses was that specific parts of the cerebellum are crucial for timing actions in relation to predictable sensory events in the cart-pole balancing task. Skilled motor behaviour, as necessary in cart-pole balancing, is suggested to rely on accurate predictive models of both our own body and tools we interact with^31,42,43^. Following this consideration, internal forward models, which represent the new tools or objects, have to be acquired during the skill acquisition process^43^. Coherently, we have previously shown^26^ that performing the cart-pole balancing task facilitates the ability to extrapolate the pole motion in a pure perception task, supporting the notion of an acquired forward model of object dynamics. We have previously also shown that predictive action timing and cart-pole balancing performance are tightly coupled^27^. Here, we verified this result and found that the cart-pole balancing performance (rnTL) is significantly correlated (r=0.71, p<0.001, N=38) with the action timing (AT) at the end of the training period. Moreover, we found no significant difference in the learning rate of action timing between the unaffected patients and the control group, while the learning rate of the affected cerebellar group was significantly different from both of the other groups (see Supplementary Material S2). This result suggests that control subjects and unaffected cerebellar subjects were able to acquire the necessary control knowledge to execute their actions predictively with respect to the pole movement. Affected cerebellar subjects, in contrast, were not able to learn to execute their actions more predictively and thus did not manage to increase the gravity over a certain level (**Fig 4** B).

These results are in line with earlier studies, showing the impairment of cerebellar patients in predictive motor timing in visual interception tasks^44–46^.

### Influence of the cerebellum on hand-independent representations for manipulating dynamic objects

Internal forward models, which predict the dynamics of the cart-pole system and which can be used for pure perceptual tasks^26^, could be at least partly hand-independent and thus should facilitate the transfer to the other hand. This hypothesis is supported for example by the study of Morton et al.^47^, in which inter-limb generalization of adaptation during ball catching was observed. Thus, it was suggested that partial inter-limb transfer is due to an internal representation of ball momentum, which is used for predictively controlling any of the hands to catch the ball. Furthermore, these authors have suggested the cerebellum as location for such a representation based on their observation that individuals with cerebellar damage have an impaired ability to adapt to a novel ball weight during catching^48^.

Both, the (1) occurrence of transfer as well as the (2) impaired transfer due to cerebellar lesions is reflected in our data. (1) The identified correlations between action timing of both hands for controls (rho=0.74, p<0.001, N=19) as well for cerebellar subjects (rho=0.54, p=0.02, N=18, **Fig 5** B) indicate that subjects were able to transfer this knowledge to the non-dominant hand on the fifth day (**Fig 4** A). This result suggests that the learned representation for predicting the cart-pole system’s behaviour and the therewith-associated ability to time actions in relation to the system’s state is not exclusively bound to the hand, which was used for acquiring the skill. (2) On the other hand, we have shown that the transfer from the dominant to the non-dominant hand was for cerebellar subjects less complete than for control subjects (**Fig 3** F), which confirms the initial hypothesis of a negative influence of impaired internal models predicting the dynamics of the cart-pole system.

#### Focal lesions associated with impaired control and learning performance

The action timing in the cart-pole balancing task measures the ability to time finger movements in relation to the predictable pole movement. Accordingly, we found impairments to be associated with lesions in the intermediate and lateral areas of lobule V and VI. The lobules V and VI are known to be involved in general hand and finger control^49,50^, of which in particular lobule VI is involved in complex movements. Especially in complex movement sequences both hemispheres are activated^33^. These considerations strongly suggest that the necessary fine-motoric finger movements in combination with the complexity of the cart-pole balancing task require bilateral cerebellar activity and, thus, legitimates our strategy to flip all lesions to the right hemisphere^51^.

Lesions in these areas are commonly also associated with decreased accuracy of finger movements^52,53^. The impaired control capabilities can therefore, firstly, be explained by the inability to execute finger movements accurately. Secondly, the lobules V and VI together with the dentate nucleus are also important for predictive action timing in target interception^46,54^. In the study by Bares et al. ^46^, successful performance was associated with increased activity in the cerebellum (right dentate nucleus as well as lobules V and VI) for healthy and cerebellar subjects.

Bilateral cerebellar activity during learning to use a new tool was also reported by Imamizu et al.^55^ and was interpreted as indication that activated regions might acquire internal models for cognitive function independent of the ipsilateral correspondence between the motor apparatus and the cerebellum. In combination with previous work, these authors suggested that internal models for the motor apparatus are present in phylogenetically older parts of the cerebellum (such as the ventral paraflocculus, vermis and intermediate parts), whereas internal models of objects and tools in the external world seem to reside in newer parts located in the hemispheres (see Imamizu et al.^55^ for a more detailed discussion).

This theory fits, on the one hand, perfectly to our observation that decreased performance in learning to balance the inverted pendulum is associated with lesions in the intermediate and lateral parts of lobules V and VI. On the other hand, it supports our hypothesis that internal models, which predict the dynamics of the manipulated object, are stored are in the cerebellum. These internal models might be at least partly stored in both hemispheres independently from the training hand and are engaged in the transfer of control knowledge to the other hand.

### Influence of the cerebellum on reward-based learning processes

While error-based or supervised learning have been suggested to rely on the cerebellum^9,56,57^, reward-based learning mechanisms are not primarily proposed to be dependent on the cerebellum but on the basal ganglia^58^. Accordingly, it was hypothesized that cerebellar patients could potentially use reward-based learning as an alternative mechanism^21^. However, increasing evidence due to recent research suggests that reward-based learning may be also supported by the cerebellum^22,59,60^.

Thus, cerebellar dysfunction might influence reward-based learning besides the previously discussed negative influence of motor variability^22,61^. For instance, BOLD response in the cerebellum has been found to be correlated with reward prediction error^62,63^. In addition, a recent neurophysiological study showed that granule cells in lobule VI of mice encode not only movement but also the expectation of reward^64^. Together with former functional-MRI studies, which indicate that lobule VI encodes sensory prediction error^65^, these results could suggest that these cerebellar regions form predictions in both mechanisms of learning: sensory predictions in the case of error-based learning and predictions of future reward for reward-based learning.

However, the above-described potential influences of cerebellar lesions on reward-based learning have to remain speculative because there is no possibility for us to distinguish such processes based on the here reported behavioural or MRT analysis.

### Interacting learning mechanisms in skill acquisition

Although the cart-pole task is a classical reinforcement learning task^23^ with only the reward of success or failure as available feedback, successful control mechanisms are suggested to involve a forward model of the cart-pole dynamics^24,25^. For our experiment this hypothesis implies that forward models have to be formed initially and recalibrated later on to different gravity levels ^43^ by reducing the error between predicted and actual cart-pole system behavior resulting in error-based learning. Consequently, acquiring the cart-pole balancing skill would require interacting learning processes: (1) acquiring and recalibrating a forward model of cart-pole dynamics and (2) reward-based learning of the valuable actions and/or system states. The interaction of both mechanisms together has previously been referred to as model-based action selection^66,67^.

Our findings of increased action variability, decreased predictive timing and the impaired transfer to the other hand are consistent with the hypothesis that impaired or poorly calibrated internal forward models represented in the cerebellum influence the skill acquisition process negatively. This hypothesis is further supported by the identified cerebellar areas, which were associated with impaired learning performance in the cart-pole task and overlap with areas commonly active in visuomotor adaptations tasks and which are suggested to represent internal forward models of the motor apparatus as well as of external objects and tools (see for example Imamizu et al.^55^).

Following this hypothesis, impaired cerebellar error-based learning such as in cerebellar patients could influence reward-based learning in motor tasks negatively by leading to poorly calibrated forward models, increased motor variability^22^, decreased predictive motor control capabilities and, finally, to less success in the task.

Another potentially relevant interaction of learning mechanisms was recently described by Wong et al. ^68^. They reported that impairments of cerebellar patients in learning rule-based strategies are caused by dysfunctional sensory predictions (see also Donchin et al. ^69^ for discussion). The development, expression and use of strategies have been suggested to play a pivotal role in reinforcement-based motor learning ^70,71^. Thus, also in our cart-pole balancing task, dysfunctional sensory predictions could influence the development of rule-based strategies negatively. Future research has to examine the interactions between these learning mechanisms further and explore training methods, which enable cerebellar patients to utilize their remaining motor learning capabilities.

## Conclusion

We have shown that even cerebellar subjects with only very mild to moderate ataxia symptoms can show significant deficits in the in learning to control the virtual cart-pole system. These deficits depend on the lesioned regions in the cerebellum. Intermediate and lateral areas of the cerebellar lobules V, VI were most strongly associated with impaired performance. Intriguingly, affected cerebellar subject showed more offline than online improvements and their learning progress does not seem to be saturated after five days. Further investigation is required to examine the long-term learning capabilities of cerebellar patients as well as to disentangle the neural mechanisms of motor learning in this population.

## Acknowledgements

NL would like to thank the German National Academic Foundation for granting a doctoral student fellowship. The study was financially supported by the Centre for Integrative Neuroscience (CIN PP-2013-1). Additional support has been received from the German Research Foundation (DFG GZ: KA 1258/15-1; TI 239/16-1), European Union Seventh Framework Programme (CogIMon H2020 ICT-23-2014/644727), the Human Frontiers Science Program (HFSP RGP0036/2016), BMBF (FKZ 01GQ1704), BW Stiftung (project KONSENS NEU007/1). We acknowledge the support by the Deutsche Forschungsgemeinschaft and Open Access Publishing Fund of University of Tübingen.

## REFERENCES

1. Trouillas, P. et al. International Cooperative Ataxia Rating Scale for pharmacological assessment of the cerebellar syndrome. Journal of the Neurological Sciences 145, 205–211; 10.1016/S0022-510X(96)00231-6 (1997).

2. Donchin, O. et al. Cerebellar regions involved in adaptation to force field and visuomotor perturbation. Journal of Neurophysiology 107, 134–147; 10.1152/jn.00007.2011 (2011).

3. Martin, T. A., Keating, J. G., Goodkin, H. P., Bastian, A. J. & Thach, W. T. Throwing while looking through prisms: I. Focal olivocerebellar lesions impair adaptation. Brain 119, 1183–1198; 10.1093/brain/119.4.1183 (1996).

4. Synofzik, M., Lindner, A. & Thier, P. The Cerebellum Updates Predictions about the Visual Consequences of One’s Behavior. Current Biology 18, 814–818; 10.1016/j.cub.2008.04.071 (2008).

5. Maschke, M. Hereditary Cerebellar Ataxia Progressively Impairs Force Adaptation During Goal-Directed Arm Movements. Journal of Neurophysiology 91, 230–238; 10.1152/jn.00557.2003 (2003).

6. Smith, M. A. & Shadmehr, R. Intact Ability to Learn Internal Models of Arm Dynamics in Huntington’s Disease But Not Cerebellar Degeneration. Journal of Neurophysiology 93, 2809–2821; 10.1152/jn.00943.2004 (2005).

7. Morton, S. M. & Bastian, A. J. Cerebellar Contributions to Locomotor Adaptations during Splitbelt Treadmill Walking. Journal of Neuroscience 26, 9107–9116; 10.1523/JNEUROSCI.2622-06.2006 (2006).

8. Bastian, A. J. Learning to predict the future: the cerebellum adapts feedforward movement control. Current Opinion in Neurobiology 16, 645–649; 10.1016/j.conb.2006.08.016 (2006).

9. Bastian, A. J. Moving, sensing and learning with cerebellar damage. Current Opinion in Neurobiology 21, 596–601; 10.1016/j.conb.2011.06.007 (2011).

10. Blakemore, S.-J., Frith, C. D. & Wolpert, D. M. The cerebellum is involved in predicting the sensory consequences of action. Neuroreport 12, 1879–1884; 10.1097/00001756-200107030-00023 (2001).

11. Bastian, A. J., Martin, T. A., Keating, J. G. & Thach, W. T. Cerebellar ataxia: abnormal control of interaction torques across multiple joints. Journal of Neurophysiology 76, 492–509 (1996).

12. Kurtzer, I. et al. Cerebellar damage diminishes long-latency responses to multijoint perturbations. Journal of Neurophysiology 109, 2228–2241; 10.1152/jn.00145.2012 (2013).

13. Flanagan, J. R., Vetter, P., Johansson, R. S. & Wolpert, D. M. Prediction Precedes Control in Motor Learning. Current Biology 13, 146–150; 10.1016/S0960-9822(03)00007-1 (2003).

14. Wolpert, D. M., Ghahramani, Z. & Flanagan, J. R. Perspectives and problems in motor learning. Trends in Cognitive Sciences 5, 487–494; 10.1016/S1364-6613(00)01773-3 (2001).

15. Ilg, W. et al. Intensive coordinative training improves motor performance in degenerative cerebellar disease. Neurology 73, 1823–1830; 10.1212/WNL.0b013e3181c33adf (2009).

16. Ilg, W. et al. Video game-based coordinative training improves ataxia in children with degenerative ataxia. Neurology 79, 2056–2060; 10.1212/WNL.0b013e3182749e67 (2012).

17. Keller, J. L. & Bastian, A. J. A home balance exercise program improves walking in people with cerebellar ataxia. Neurorehabilitation and Neural Repair 28, 770–778; 10.1177/1545968314522350 (2014).

18. Ilg, W. et al. Consensus paper. Management of degenerative cerebellar disorders. Cerebellum 13, 248–268; 10.1007/s12311-013-0531-6 (2014).

19. Morton, S. M. & Bastian, A. J. Can rehabilitation help ataxia? Neurology 73, 1818–1819; 10.1212/WNL.0b013e3181c33b21 (2009).

20. Doya, K. Complementary roles of basal ganglia and cerebellum in learning and motor control. Current Opinion in Neurobiology 10, 732–739; 10.1016/S0959-4388(00)00153-7 (2000).

21. Criscimagna-Hemminger, S. E., Bastian, A. J. & Shadmehr, R. Size of Error Affects Cerebellar Contributions to Motor Learning. Journal of Neurophysiology 103, 2275–2284; 10.1152/jn.00822.2009 (2010).

22. Therrien, A. S., Wolpert, D. M. & Bastian, A. J. Effective reinforcement learning following cerebellar damage requires a balance between exploration and motor noise. Brain 139, 101–114; 10.1093/brain/awv329 (2016).

23. Barto, A. G., Sutton, R. S. & Anderson, C. W. Neuronlike adaptive elements that can solve difficult learning control problems. IEEE Transactions on Systems, Man, and Cybernetics 13, 834–846; 10.1109/TSMC.1983.6313077 (1983).

24. Mehta, B. & Schaal, S. Forward Models in Visuomotor Control. Journal of Neurophysiology 88, 942–953 (2002).

25. Lee, K.-Y., O’Dwyer, N., Halaki, M. & Smith, R. Perceptual and motor learning underlies human stick-balancing skill. Journal of Neurophysiology 113, 156–171; 10.1152/jn.00538.2013 (2015).

26. Ludolph, N., Plöger, J., Giese, M. A. & Ilg, W. Motor expertise facilitates the accuracy of state extrapolation in perception. PLoS ONE 12, e0187666; 10.1371/journal.pone.0187666 (2017).

27. Ludolph, N., Giese, M. A. & Ilg, W. Interacting Learning Processes during Skill Acquisition. Learning to control with gradually changing system dynamics. Scientific reports 7, 13191; 10.1038/s41598-017-13510-0 (2017).

28. Dayan, E. & Cohen, L. G. Neuroplasticity Subserving Motor Skill Learning. Neuron 72, 443–454; 10.1016/j.neuron.2011.10.008 (2011).

29. Reis, J. et al. Noninvasive cortical stimulation enhances motor skill acquisition over multiple days through an effect on consolidation. Proceedings of the National Academy of Sciences 106, 1590–1595; 10.1073/pnas.0805413106 (2009).

30. Miall, R. C. & Galea, J. Cerebellar damage limits reinforcement learning. Brain 139, 4–7; 10.1093/brain/awv343 (2016).

31. Imamizu, H. Modular organization of internal models of tools in the human cerebellum. Proceedings of the National Academy of Sciences 100, 5461–5466; 10.1073/pnas.0835746100 (2003).

32. Oldfield, R. C. The assessment and analysis of handedness. The Edinburgh inventory. Neuropsychologia 9, 97–113; 10.1016/0028-3932(71)90067-4 (1971).

33. Schlerf, J. E., Verstynen, T. D., Ivry, R. B. & Spencer, R. M. C. Evidence of a novel somatopic map in the human neocerebellum during complex actions. Journal of Neurophysiology 103, 3330–3336; 10.1152/jn.01117.2009 (2010).

34. Iglewicz, B. & Hoaglin, D. C. How to Detect and Handle Outliers (ASQC Quality Press, Milwaukee, 1993).

35. Diedrichsen, J. A spatially unbiased atlas template of the human cerebellum. NeuroImage 33, 127–138; 10.1016/j.neuroimage.2006.05.056 (2006).

36. Karnath, H.-O., Himmelbach, M. & Rorden, C. The subcortical anatomy of human spatial neglect. Putamen, caudate nucleus and pulvinar. Brain 125, 350–360; 10.1093/brain/awf032 (2002).

37. Rorden, C., Karnath, H.-O. & Bonilha, L. Improving Lesion-Symptom Mapping. Journal of Cognitive Neuroscience 19, 1081–1088; 10.1162/jocn.2007.19.7.1081 (2007).

38. Cantarero, G. et al. Cerebellar direct current stimulation enhances on-line motor skill acquisition through an effect on accuracy. The Journal of Neuroscience 35, 3285–3290; 10.1523/JNEUROSCI.2885-14.2015 (2015).

39. Galea, J. M., Vazquez, A., Pasricha, N., Orban de Xivry, J.-J. & Celnik, P. A. Dissociating the Roles of the Cerebellum and Motor Cortex during Adaptive Learning: The Motor Cortex Retains What the Cerebellum Learns. Cerebral Cortex 21, 1761–1770; 10.1093/cercor/bhq246 (2011).

40. Gerwig, M. et al. Evaluation of multiple-session delay eyeblink conditioning comparing patients with focal cerebellar lesions and cerebellar degeneration. Behavioural brain research 212, 143–151; 10.1016/j.bbr.2010.04.007 (2010).

41. Chu, V. W. T., Sternad, D. & Sanger, T. D. Healthy and dystonic children compensate for changes in motor variability. Journal of Neurophysiology 109, 2169–2178; 10.1152/jn.00908.2012 (2013).

42. Dingwell, J. B., Mah, C. D. & Mussa-Ivaldi, F. A. Manipulating Objects With Internal Degrees of Freedom: Evidence for Model-Based Control. Journal of Neurophysiology 88, 222–235 (2002).

43. Wolpert, D. M. & Flanagan, J. R. Motor prediction. Current Biology 11, R729–R732; 10.1016/S0960-9822(01)00432-8 (2001).

44. Broersen, R. et al. Impaired Spatio-Temporal Predictive Motor Timing Associated with Spinocerebellar Ataxia Type 6. PLoS ONE 11, e0162042; 10.1371/journal.pone.0162042 (2016).

45. Bares, M. et al. Impaired predictive motor timing in patients with cerebellar disorders. Experimental Brain Research 180, 355–365; 10.1007/s00221-007-0857-8 (2007).

46. Bares, M. et al. The neural substrate of predictive motor timing in spinocerebellar ataxia. Cerebellum 10, 233–244; 10.1007/s12311-010-0237-y (2011).

47. Morton, S. M., Lang, C. E. & Bastian, A. J. Inter- and intra-limb generalization of adaptation during catching. Experimental Brain Research 141, 438–445; 10.1007/s002210100889 (2001).

48. Lang, C. E. & Bastian, A. J. Cerebellar subjects show impaired adaptation of anticipatory EMG during catching. Journal of Neurophysiology 82, 2108–2119 (1999).

49. Grodd, W., Hülsmann, E., Lotze, M., Wildgruber, D. & Erb, M. Sensorimotor mapping of the human cerebellum: fMRI evidence of somatotopic organization. Hum. Brain Mapp. 13, 55–73; 10.1002/hbm.1025 (2001).

50. Diedrichsen, J. & Zotow, E. Surface-Based Display of Volume-Averaged Cerebellar Imaging Data. PLoS ONE 10, e0133402; 10.1371/journal.pone.0133402 (2015).

51. Timmann, D. et al. Current advances in lesion-symptom mapping of the human cerebellum. Neuroscience 162, 836–851; 10.1016/j.neuroscience.2009.01.040 (2009).

52. Glickstein, M., Waller, J., Baizer, J., Brown, B. & Timmann, D. Cerebellum lesions and finger use. The Cerebellum 4, 189–197; 10.1080/14734220500201627 (2005).

53. Brandauer, B. et al. Impaired and preserved aspects of independent finger control in patients with cerebellar damage. Journal of Neurophysiology 107, 1080–1093; 10.1152/jn.00142.2011 (2012).

54. Husárová, I. et al. Functional imaging of the cerebellum and basal ganglia during predictive motor timing in early Parkinson’s disease. Journal of neuroimaging: official journal of the American Society of Neuroimaging 24, 45–53; 10.1111/j.1552-6569.2011.00663.x (2014).

55. Imamizu, H. et al. Human cerebellar activity reflecting an acquired internal model of a new tool. Nature 403, 192–195; 10.1038/35003194 (2000).

56. Shadmehr, R., Smith, M. A. & Krakauer, J. W. Error Correction, Sensory Prediction, and Adaptation in Motor Control. Annual Review of Neuroscience 33, 89–108; 10.1146/annurev-neuro-060909-153135 (2010).

57. Wolpert, D. M., Diedrichsen, J. & Flanagan, J. R. Principles of sensorimotor learning. Nature Reviews Neuroscience 12, 739–751; 10.1038/nrn3112 (2011).

58. Schultz, W., Dayan, P. & Montague, P. R. A Neural Substrate of Prediction and Reward. Science 275, 1593–1599; 10.1126/science.275.5306.1593 (1997).

59. Haith, A. M. & Krakauer, J. W. in Progress in Motor Control, edited by M. J. Richardson, M. A. Riley & K. Shockley (Springer New York 2013), pp. 1–21.

60. Taylor, J. A. & Ivry, R. B. Cerebellar and prefrontal cortex contributions to adaptation, strategies, and reinforcement learning. Progress in Brain Research 210, 217–253; 10.1016/B978-0-444-63356-9.00009-1 (2014).

61. Therrien, A. S., Wolpert, D. M. & Bastian, A. J. Increasing Motor Noise Impairs Reinforcement Learning in Healthy Individuals. eNeuro 5; 10.1523/ENEURO.0050-18.2018 (2018).

62. O’Doherty, J. P., Dayan, P., Friston, K., Critchley, H. & Dolan, R. J. Temporal Difference Models and Reward-Related Learning in the Human Brain. Neuron 38, 329–337; 10.1016/S0896-6273(03)00169-7 (2003).

63. Seymour, B. et al. Temporal difference models describe higher-order learning in humans. Nature 429, 664–667; 10.1038/nature02581 (2004).

64. Wagner, M. J., Kim, T. H., Savall, J., Schnitzer, M. J. & Luo, L. Cerebellar granule cells encode the expectation of reward. Nature 544, 96–100; 10.1038/nature21726 (2017).

65. Schlerf, J., Ivry, R. B. & Diedrichsen, J. Encoding of sensory prediction errors in the human cerebellum. The Journal of neuroscience: the official journal of the Society for Neuroscience 32, 4913–4922; 10.1523/JNEUROSCI.4504-11.2012 (2012).

66. Doya, K. What are the computations of the cerebellum, the basal ganglia and the cerebral cortex? Neural Networks 12, 961–974; 10.1016/S0893-6080(99)00046-5 (1999).

67. Caligiore, D. et al. Consensus Paper. Towards a Systems-Level View of Cerebellar Function: the Interplay Between Cerebellum, Basal Ganglia, and Cortex. Cerebellum 16, 203–229; 10.1007/s12311-016-0763-3 (2017).

68. Wong, A. L., Marvel, C. L., Taylor, J. A. & Krakauer, J. W. Can patients with cerebellar disease switch learning mechanisms to reduce their adaptation deficits? Brain 142, 662–673; 10.1093/brain/awy334 (2019).

69. Donchin, O. & Timmann, D. How to help cerebellar patients make the most of their remaining learning capacities. Brain 142, 492–495; 10.1093/brain/awz020 (2019).

70. Holland, P., Codol, O. & Galea, J. M. Contribution of explicit processes to reinforcement-based motor learning. Journal of Neurophysiology 119, 2241–2255; 10.1152/jn.00901.2017 (2018).

71. Codol, O., Holland, P. J. & Galea, J. M. The relationship between reinforcement and explicit control during visuomotor adaptation. Scientific reports 8, 9121; 10.1038/s41598-018-27378-1 (2018).

72. Brainard, D. H. The Psychophysics Toolbox. Spatial Vision 10, 433–436; 10.1163/156856897X00357 (1997).

73. Pelli, D. G. The VideoToolbox software for visual psychophysics: transforming numbers into movies. Spatial Vision 10, 437–442; 10.1163/156856897X00366 (1997).

74. Kleiner, M., Brainard, D. H. & Pelli, D. G. in European Conference on Visual Perception (2007).

75. Diedrichsen, J., Balsters, J. H., Flavell, J., Cussans, E. & Ramnani, N. A probabilistic MR atlas of the human cerebellum. NeuroImage 46, 39–46; 10.1016/j.neuroimage.2009.01.045 (2009).

76. Schmahmann, J. D. et al. Three-dimensional MRI atlas of the human cerebellum in proportional stereotaxic space. NeuroImage 10, 233–260; 10.1006/nimg.1999.0459 (1999).

