## Supplementary Material for "Cerebellar involvement in learning to balance a cart-pole system"

**Acknowledgements**
NL would like to thank the German National Academic Foundation for granting a doctoral student fellowship. The study was financially supported by the Centre for Integrative Neuroscience (CIN PP-2013-1). Additional support has been received from the German Research Foundation (DFG GZ: KA 1258/15-1; TI 239/16-1), European Union Seventh Framework Programme (CogIMon H2020 ICT-23-2014/644727), the Human Frontiers Science Program (HFSP RGP0036/2016), BMBF (FKZ 01GQ1704), BW Stiftung (project KONSENS NEU007/1). We acknowledge the support by the Deutsche Forschungsgemeinschaft and Open Access Publishing Fund of University of Tübingen.

### S1 – TECHNICAL DETAILS OF THE EXPERIMENTAL SETUP

The cart-pole balancing task was implemented using Psychtoolbox [1–3] for MATLAB® (The MathWorks, Inc.) on a laptop (Dell Precision M4200, 15.6" screen, 60Hz refresh rate). We used a SpaceMouse® Pro (3Dconnexion) as input device (main text: Figure 1 B). This input device is similar to a joystick but it can measure six degrees of freedom (DOF) including the left-right translation. The device's knob can be displaced by 1.5mm to the left and right while exerting a force of 7.4 N at full lateral displacement back to the centre position. The measured displacement was used to scale the virtual force in the cart-pole balancing task. The haptic input device's orientation was aligned with the laptop monitor such that the lateral knob movement was in correspondence with the cart movement in the cart-pole balancing task. For the cart-pole balancing task, we recorded for every trial the cart-pole state trajectory, current gravity level and the applied virtual force over time. All participants were instructed to sit comfortable in work posture in front of the monitor. For the home-based sessions, subjects were asked to setup the laptop and input device in the same way as in our laboratory.

#### Details of the virtual cart-pole system

The virtual cart-pole system consists of a cart to which a one-meter long pole is attached (6 cm on the screen). The masses of the pole and cart were set to 0.08 kg and 0.4 kg respectively. We did not simulate friction. Magnitude and direction of the virtual forces were calculated based on the input device's measurement (linear scaling, 0-4N, see Experimental Setup).

### S2 – MODELLING DETAILS

In order to compare the decrease in rnTL between groups, we fitted a non-linear regression model, using MATLABs Statistics and Machine Learning Toolbox™ (The Mathworks, Inc.). Our model (eq1) is a mixture of three exponential functions, one for each subgroup (affected, unaffected cerebellar and control subjects) multiplied by the respective indicator variable ($I_A, I_U$ and $I_C$), where $rnTL$ denotes the reciprocal normalized trial length and $t$ the time interacted with the cart-pole system since the beginning of the experiment. In the exponential function, the parameter $a_g$ corresponds to the final rnTL that is approached, $b_g$ represents the magnitude of learning and $c_g$ is the exponential rate at which the learning takes place. The last two parameters in combination can be interpreted as *learning rate*.

$$rnTL \sim \sum_{g\in[C,A,U]} I_g \left(a_g + b_g \, exp(-c_g \, t)\right) \qquad \text{(eq1)}$$

We accounted for outliers by weighting residuals according to the bisquare method. In order to compare the estimated coefficients of the three groups we used linear hypothesis testing and corrected for multiple comparisons (N=3) using the Bonferroni method.

#### Coefficients of fitted non-linear regression model (rnTL)

We examined the coefficients of the fitted model for learning in the different subgroups (affected, unaffected cerebellar and control subjects) using linear hypothesis testing.

| | | Estimate | SE | tStat | pValue |
|---|---|---|---|---|---|
| Control | $a_C$ | 0.15428 | 0.0039493 | 39.065 | < 0.001 |
| | $b_C$ | 0.17623 | 0.0082169 | 21.447 | < 0.001 |
| | $c_C$ | 0.049483 | 0.0052612 | 9.4053 | < 0.001 |
| Cerebellar Affected | $a_A$ | 0.42226 | 0.0030912 | 136.6 | < 0.001 |
| | $b_A$ | 0.6616 | 0.07001 | 9.4501 | < 0.001 |
| | $c_A$ | 19.993 | 6.7562 | 2.9593 | 0.0030891 |
| Cerebellar Unaffected | $a_U$ | 0.22162 | 0.0037323 | 59.378 | < 0.001 |
| | $b_U$ | 0.19394 | 0.0090541 | 21.42 | < 0.001 |
| | $c_U$ | 0.055173 | 0.005535 | 9.9682 | < 0.001 |

In linear hypothesis testing the null hypothesis $H_0: H\beta = c$ is statistically tested where $\beta$ is the parameter vector, H is used to specify the linear restrictions and c is a constant vector expressing the target value. Specifically, we tested on the difference between the parameters which represent the rate of learning ($b_g$ and $c_g$ in combination, or solely $c_g$). Given the parameter vector $\beta = (a_C, b_C, c_C, a_A, b_A, c_A, a_U, b_U, c_U)$ from the nonlinear regression model fit, we performed the tests using MATLAB's build-in function for linear hypothesis testing on nonlinear regression model coefficients (coefTest). The uncorrected results are reported in the table below. Correction for multiple comparisons was performed for reporting in the main text using Bonferroni method.

| Linear hypothesis test | | F | DoF | pValue |
|---|---|---|---|---|
| Test | $(b_C, c_C)$ vs $(b_A, c_A)$ | 25.1973 | 2 | < 0.001 |
| | $(b_C, c_C)$ vs $(b_U, c_U)$ | 1.0751 | 2 | 0.3412 |
| | $(b_A, c_A)$ vs $(b_U, c_U)$ | 23.5203 | 2 | < 0.001 |
| Test | $(c_C)$ vs $(c_A)$ | 8.7139 | 1 | 0.003 |
| | $(c_C)$ vs $(c_U)$ | 0.5552 | 1 | 0.46 |
| | $(c_A)$ vs $(c_U)$ | 8.7089 | 1 | 0.003 |

**Coefficients of fitted linear regression model (AT, subgroups)**

| | Estimate | SE | tStat | pValue |
|---|---|---|---|---|
| 1:CerebellarAffected | -17.519 | 8.7606 | -1.9998 | 0.045865 |
| 1:CerebellarUnaffected | -67.358 | 5.249 | -12.832 | 0 |
| 1:Control | -70.671 | 4.4941 | -15.725 | 0 |
| Time:CerebellarAffected | 0.24756 | 0.14455 | 1.7126 | 0.087183 |
| Time:CerebellarUnaffected | -0.18905 | 0.087127 | -2.1698 | 0.030316 |
| Time:Control | -0.31657 | 0.074155 | -4.269 | 0.000022018 |

| Linear hypothesis test | F | DoF | pValue |
|---|---|---|---|
| Time:Control vs Time:CerebellarAffected | 12.0570 | 1 | <0.001 |
| Time:Control vs Time:CerebellarUnaffected | 1.2422 | 1 | 0.27 |
| Time:CerebellarAffected vs Time:CerebellarUnaffected | 6.6918 | 1 | 0.01 |

### S3 – DETAILS ABOUT THE LESION SYMPTOM MAPPING

#### Technical details

A 3D sagittal volume of the entire brain was made using a T1-weighted, magnetization-prepared rapid acquisition gradient-echo (MPRAGE), sequence (192 sagittal slices, TR=2500ms, TE=4.38ms, TI=1100ms, acquisition matrix 256x256, pMRI GRAPPA R=2, TA=5:08min, 1mm slice thickness; non-interpolated isotropic voxel size 1.0mm). In addition, three-dimensional fluid attenuation inversion recovery (FLAIR) images were acquired (256 transversal slices, TR=5000ms, TE=395ms, TI=1800ms, turbo factor 284, acquisition matrix 256x256, pMRI GRAPPA R=3 TA=6:22min, 1mm slice thickness; non-interpolated isotropic voxel size 1.0mm). T1-weighted and FLAIR images were examined by an experienced neuroradiologist (S.G.). For none of the cerebellar subjects extracerebellar pathology was revealed. In one cerebellar subject (cp10 in main text Table 1), MR images could not be acquired because of a recently implanted contraceptive intrauterine device which was not MRI safe. We instead used T1-weighted and FLAIR images which had been acquired in a previous study using a 1.5T MRI scanner [4]. Repeated MRI scans over the previous years showed that the size of the chronic surgical lesion has remained constant.

#### Additional Lesion symptom Analysis

As part of the lesion symptom analysis, we performed a subtraction analysis [5] and Liebermeister’s tests. For performing the subtraction analysis or Liebermeister’s test, the behaviour of each subject has first to be classified as impaired or unimpaired.

The subtraction analysis can yield positive and negative values because it simply subtracts two relative frequencies. It yields a positive value for voxels that are more frequently damaged if the behaviour is impaired than if it is unimpaired, i.e. it expresses the consistency in percent. For example, highest consistency (100%) would be achieved if the voxel is only damaged if the behaviour is impaired but not if the behaviour is unimpaired. Lowest consistency (0%)

would be achieved if the voxel is equally often damaged independent of whether the behaviour is impaired or not. Finally, highest inconsistency (-100%) would be achieved if the voxel is only damaged if the behaviour is unimpaired but not if the behaviour is impaired.

The following figures (**Supplementary Figure 1**, **Supplementary Figure 3** and **Supplementary Figure 4**) show the voxel-wise lesion symptom mapping for the measure mrGL, rnTL combined hands and AT combined hands (see main text). **Supplementary Figure 2** illustrates the measure rnTL combined hands.

For performing the VLSM for each hand separately, we classified cerebellar subjects' behaviour as impaired if their single-handed performance lay outside of 99% confidence interval of control subjects' performance. **Supplementary Figure 5** shows the subtraction analysis of the measures rnTL and AT for both hands separately.

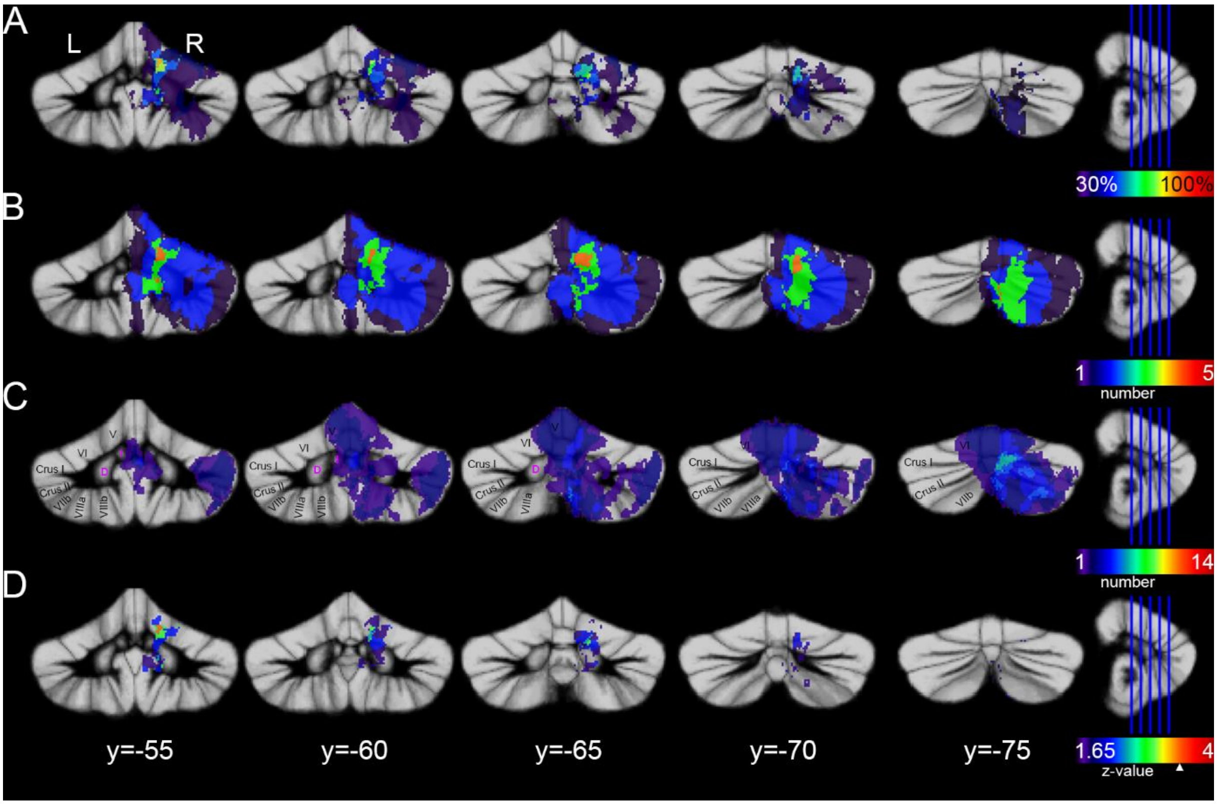


**Supplementary Figure 1. VLSM considering the maximum reached gravity (mrGL).** (A) Subtraction analysis and (D) Liebermeister test show cerebellar areas that were more likely to be lesioned in subjects with abnormally low gravity reached (mrGL<2.0). Colour in the subtraction analysis represents % consistency with a threshold of 30%. For the Liebermeister test color indicates the z-value, with a threshold of p<0.05 (uncorrected). Permutation corrected z-value (z-value=3.43) is indicated by a white triangle. Lesion-overlap images for affected (B) and unaffected (C) cerebellar subjects are shown for comparison. Results are superimposed on the maximum probability SUIT template of the cerebellum [6]. Lesions were flipped to the right side before analysis. Names of cerebellar lobules were indicated according to Schmahmann et al. [7].

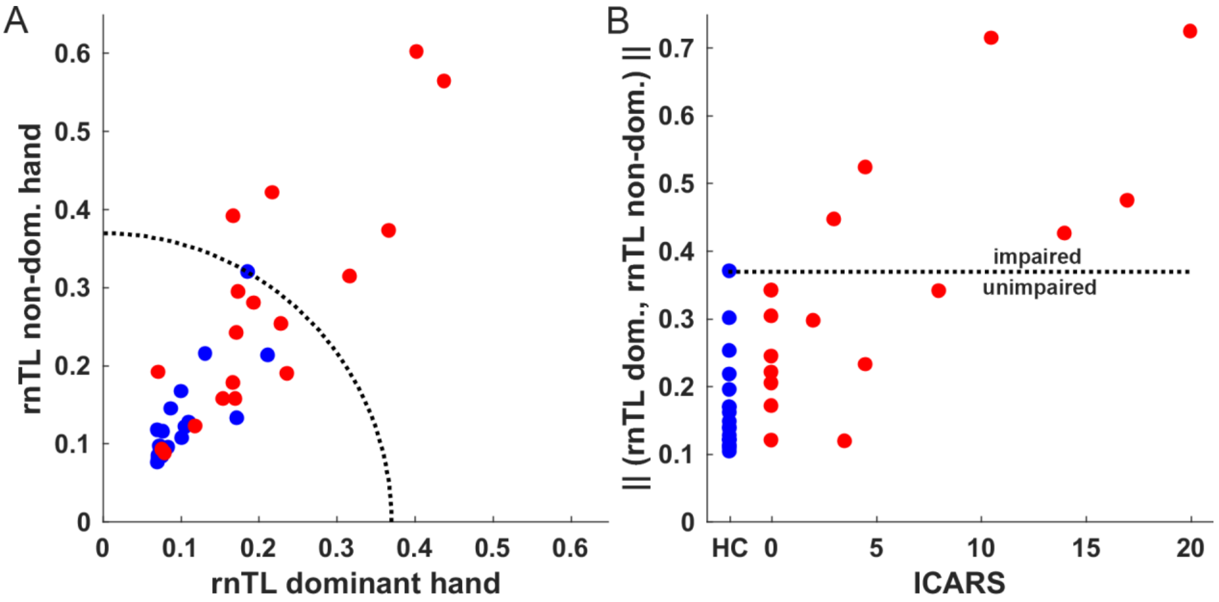


**Supplementary Figure 2. Classification of subjects' behaviors considering the reciprocal normalized trial length (rnTL).** (A) Reciprocal normalized trial length (rnTL) of the dominant versus the non-dominant hand averaged over the last 5 minutes of the respective sessions on the fifth day. (B) rnTL of the dominant and non-dominant hand combined (rnTL combined hands) as function of the ICARS score for cerebellar (red) and control subjects (blue).

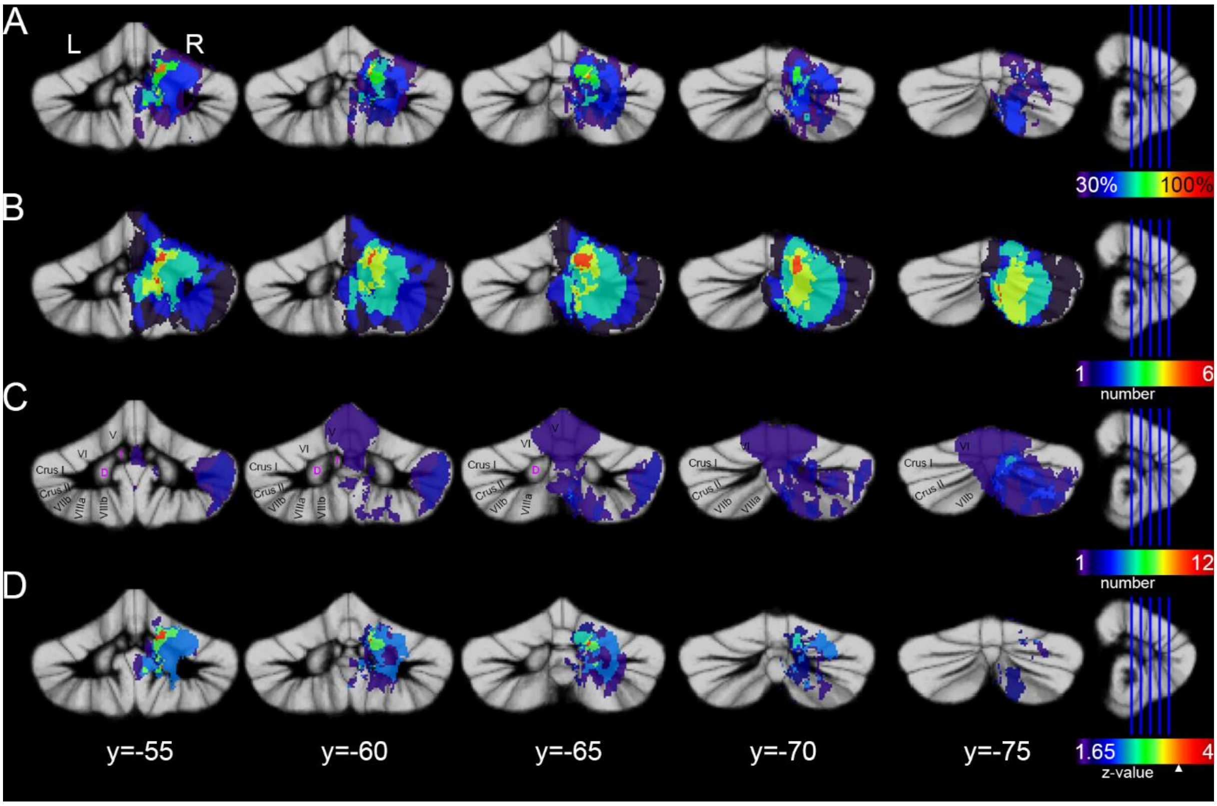


**Supplementary Figure 3. VLSM considering the reciprocal normalized trial length (rnTL) of both hands on the fifth day.** (A) Subtraction analysis and (D) Liebermeister test show cerebellar areas that were more likely to be lesioned in subjects with abnormally high rnTL. Colour in the subtraction analysis represents % consistency with a threshold of 30%. For the Liebermeister test colour indicates the z-value, with a threshold of $p<0.05$ (uncorrected). Permutation corrected z-value (z-value=3.42) is indicated by a white triangle. Lesion-overlap images for affected (B) and unaffected (C) cerebellar subjects are shown for comparison. Results are superimposed on the maximum probability SUIT template of the cerebellum [6]. Lesions were flipped to the right side before analysis. Names of cerebellar lobules were indicated according to Schmahmann et al. [7].

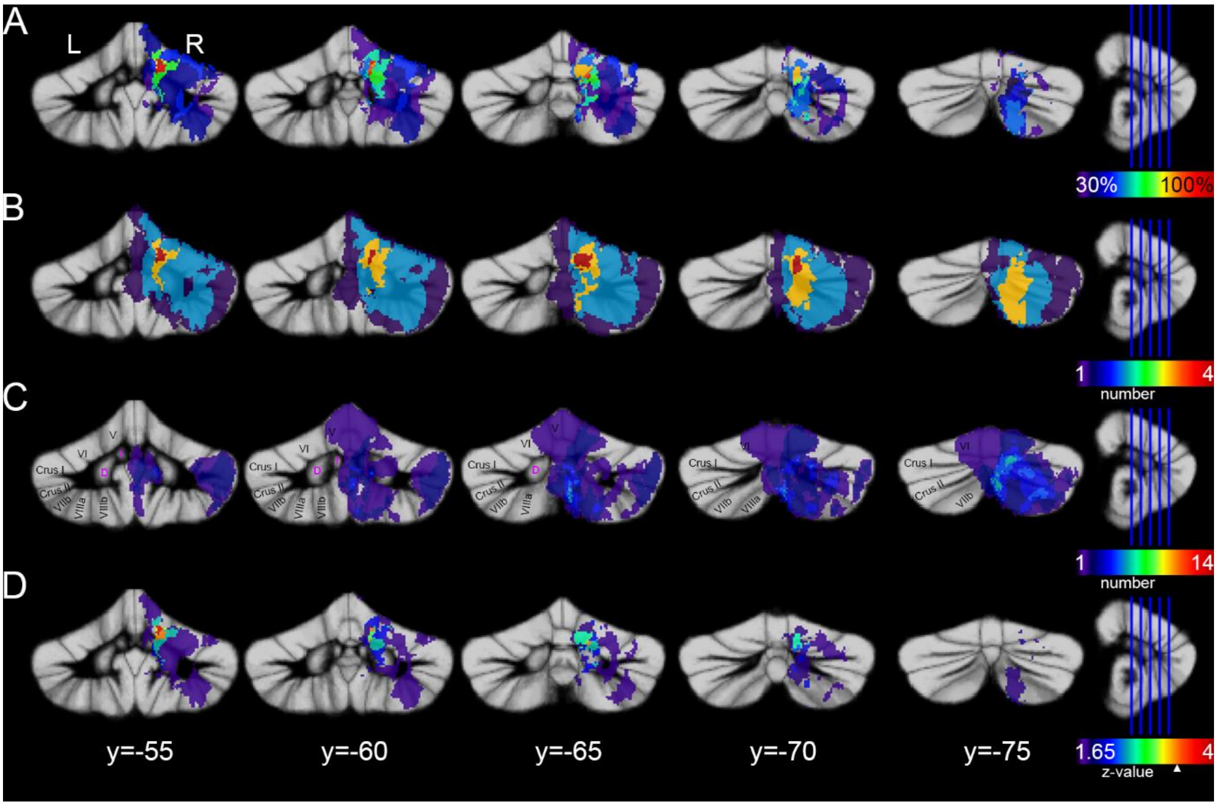


**Supplementary Figure 4. VLSM considering the action timing (AT) of both hands on the fifth day.** (A) Subtraction analysis and (D) Liebermeister test show cerebellar areas that were more likely to be lesioned in subjects with abnormally low gravity reached. Colour in the subtraction analysis represents % consistency with a threshold of 30%. For the Liebermeister test color indicates the z-value, with a threshold of $p<0.05$ (uncorrected). Permutation corrected z-value (z-value=3.36) is indicated by a white triangle. Lesion-overlap images for affected (B) and unaffected (C) cerebellar subjects are shown for comparison. Results are superimposed on the maximum probability SUIT template of the cerebellum [6]. Lesions were flipped to the right side before analysis. Names of cerebellar lobules were indicated according to Schmahmann et al. [7].

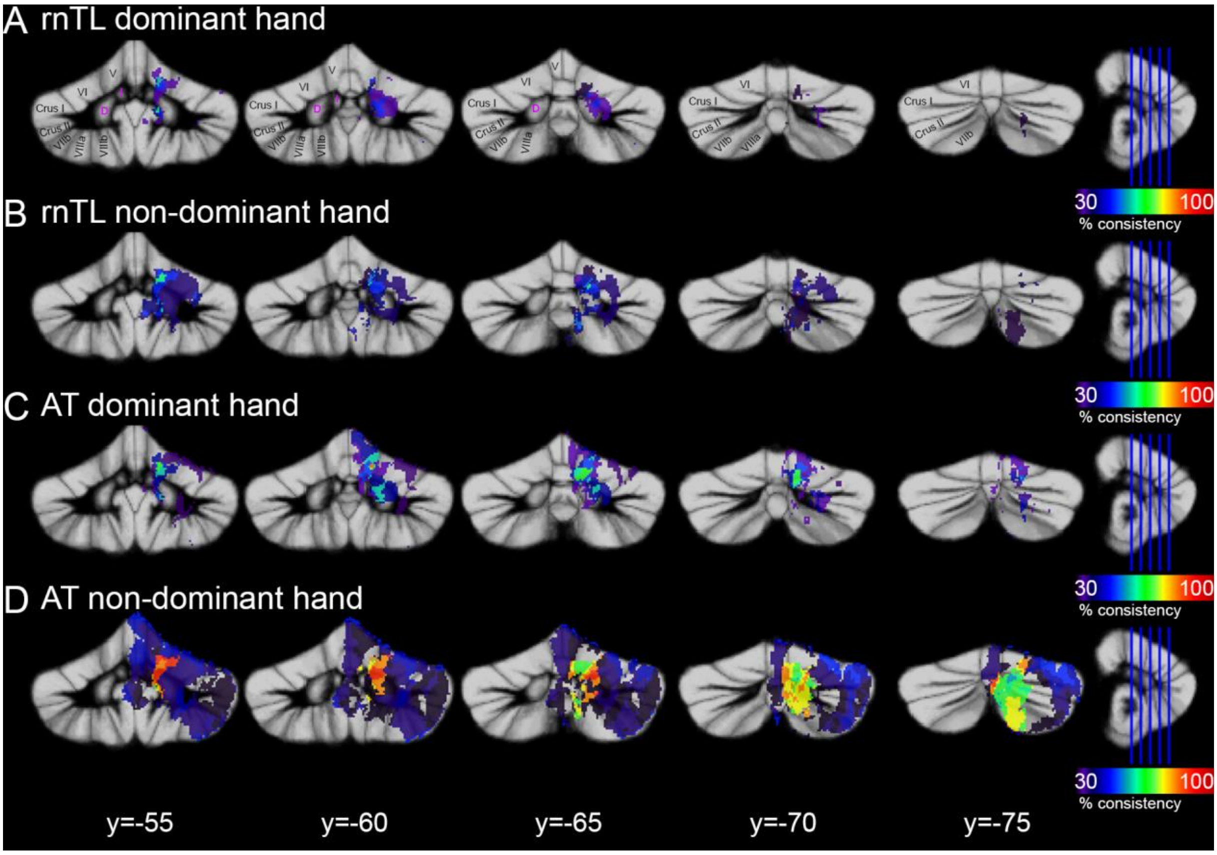


**Supplementary Figure 5. VLSM considering the reciprocal normalized trial length (rnTL) and action timing (AT) for each hand on the fifth day.** (A, B) Subtraction analysis considering the measure rnTL and (C, D) considering the measure AT for the (A, C) dominant and (B, D) non-dominant hand respectively. The colour represents the percentage consistency of impaired behaviour and lesion. Notice, that only two subjects were classified as impaired regarding the action timing using the non-dominant hand. All results are superimposed on the maximum probability SUIT template of the cerebellum [6]. Lesions were flipped to the right side before analysis. Names of cerebellar lobules were indicated according to Schmahmann et al. [7].

### REFERENCES

1. Brainard, D. H. The Psychophysics Toolbox. *Spatial Vision* **10,** 433–436; 10.1163/156856897X00357 (1997).
2. Pelli, D. G. The VideoToolbox software for visual psychophysics: transforming numbers into movies. *Spatial Vision* **10,** 437–442; 10.1163/156856897X00366 (1997).
3. Kleiner, M., Brainard, D. H. & Pelli, D. G. in *European Conference on Visual Perception* (2007).

4. Brandauer, B. *et al.* Impaired and preserved aspects of independent finger control in patients with cerebellar damage. *Journal of Neurophysiology* **107,** 1080–1093; 10.1152/jn.00142.2011 (2012).

5. Karnath, H.-O., Himmelbach, M. & Rorden, C. The subcortical anatomy of human spatial neglect. Putamen, caudate nucleus and pulvinar. *Brain* **125,** 350–360; 10.1093/brain/awf032 (2002).

6. Diedrichsen, J., Balsters, J. H., Flavell, J., Cussans, E. & Ramnani, N. A probabilistic MR atlas of the human cerebellum. *NeuroImage* **46,** 39–46; 10.1016/j.neuroimage.2009.01.045 (2009).

7. Schmahmann, J. D. *et al.* Three-dimensional MRI atlas of the human cerebellum in proportional stereotaxic space. *NeuroImage* **10,** 233–260; 10.1006/nimg.1999.0459 (1999).
